# Simultaneous estimation of gene regulatory network structure and RNA kinetics from single cell gene expression

**DOI:** 10.1101/2023.09.21.558277

**Authors:** Christopher A Jackson, Maggie Beheler-Amass, Andreas Tjärnberg, Ina Suresh, Angela Shang-mei Hickey, Richard Bonneau, David Gresham

## Abstract

Cells respond to environmental and developmental stimuli by remodeling their transcriptomes through regulation of both mRNA transcription and mRNA decay. A central goal of biology is identifying the global set of regulatory relationships between factors that control mRNA production and degradation and their target transcripts and construct a predictive model of gene expression. Regulatory relationships are typically identified using transcriptome measurements and causal inference algorithms. RNA kinetic parameters are determined experimentally by employing run-on or metabolic labeling (e.g. 4-thiouracil) methods that allow transcription and decay rates to be separately measured. Here, we develop a deep learning model, trained with single-cell RNA-seq data, that both infers causal regulatory relationships and estimates RNA kinetic parameters. The resulting *in silico* model predicts future gene expression states and can be perturbed to simulate the effect of transcription factor changes.

We acquired model training data by sequencing the transcriptomes of 175,000 individual *Saccharomyces cerevisiae* cells that were subject to an external perturbation and continuously sampled over a one hour period. The rate of change for each transcript was calculated on a per-cell basis to estimate RNA velocity. We then trained a deep learning model with transcriptome and RNA velocity data to calculate time-dependent estimates of mRNA production and decay rates. By separating RNA velocity into transcription and decay rates, we show that rapamycin treatment causes existing ribosomal protein transcripts to be rapidly destabilized, while production of new transcripts gradually slows over the course of an hour.

The neural network framework we present is designed to explicitly model causal regulatory relationships between transcription factors and their genes, and shows superior performance to existing models on the basis of recovery of known regulatory relationships. We validated the predictive power of the model by perturbing transcription factors *in silico* and comparing transcriptome-wide effects with experimental data. Our study represents the first step in constructing a complete, predictive, biophysical model of gene expression regulation.

## 1 Introduction

Gene expression regulation is a multifaceted process that commences with transcription, followed by processing, translation, and degradation of mRNA (*1*). Transcription is regulated by transcription factors (TFs), which influence and are influenced by chromatin accessibility, and control the production of new precursor mRNA (pre-mRNA). Pre-mRNA molecules are packaged into messenger ribonucleoprotein particles with RNA-binding proteins, which directly and indirectly control pre-mRNA splicing, nuclear export, translation, and mRNA degradation (*2*). mRNA decay is regulated (*3*) in response to changes in the extracellular environment (*4, 5*) and is frequently dysregulated in disease (*6*). Quantifying mRNA production and decay rates is technically challenging as it requires metabolic labeling of transcripts (*7,8*) to infer transcription and degradation from change in labeled RNA over time (*9*).

A central goal in understanding gene expression regulation is constructing gene regulatory networks (GRNs) that define the regulatory relationships between genes and the factors that control their expression. A complete GRN model should predict gene expression states over time and in response to perturbation of network components (e.g. deletion or overexpression of TFs). Many computational methods exist to infer GRNs, which are typically represented as directed graphs that connect TFs to their target genes (*10–13*). The most successful methods have a common framework; a prior knowledge GRN is generated based on existing evidence of regulatory relationships, and a regression or classification algorithm uses the prior knowledge GRN with experimentally obtained gene expression or chromatin accessibility data to identify novel high probability regulatory relationships between TFs and genes. Time series data are extremely useful for learning GRNs (*14*) as coordinated temporal dynamics are evidence of regulatory relationships (*15–17*). However, high quality time series data sets are rare as they are laborious to collect.

Single-cell RNA-sequencing (scRNA-seq) methods have greatly expanded the number of individual genome-wide observations that can be made from a cell population. Technological advances have improved sequencing depth and throughput from a handful of cells (*18*) to millions of cells (*19*), enabling single-cell analysis of dynamic, time-dependent processes like the cell cycle (*20*), and response to drug treatment (*21*). Many methods have been proposed to learn GRNs from scRNA-seq data, although the lack of a reliable gold-standard networks for most organisms has limited rigorous evaluation of these methods to synthetic benchmarks (*22*), and the optimal approach to constructing GRNs from scRNA-seq data remains an open question.

Ideally, GRNs would be defined using rates of transcription as TFs directly control the production of mRNA, but not its degradation. However, experimental transcription rate data is difficult to collect, and as a result GRNs are generally inferred based on transcript abundance data. Typically, the rate of mRNA decay is not modeled in GRNs. However, initial efforts to explicitly incorporate mRNA decay parameters as a tunable network inference model parameter improved network inference performance (*23*) suggesting that incorporation of kinetic parameters may enhance GRN reconstruction.

Many organisms splice non-coding introns from pre-mRNA (*24*), and the rate of transcriptional output can be estimated by comparing the ratio of intronic to exonic sequence reads (*25*). This approach has been applied to scRNA-seq expression data to estimate RNA velocity (*26, 27*), defined as the rate of change over time of an mRNA transcript, addressing questions regarding cell developmental trajectories (*28*) and biophysical parameter estimation (*29*). However, inherent limitations in the ability to capture intronic reads make this approach challenging (*27*), particularly with scRNA-seq methods that target the 3’ end of transcripts. The use of RNA velocity in GRN inference is still preliminary (*30, 31*) although recent work has extended the concept of RNA velocity to predictive models based on deep learning combined with ordinary differential equation models to predict gene expression at future states and to infer GRNs (*32–34*).

Here, we developed a dynamical deep learning model based on biologically informed neural networks (*35*) to simultaneously infer the structure of a GRN and estimate the rate of mRNA transcription and decay using scRNA-seq data. In the model, mRNA transcription is causally linked to regulatory TFs allowing for *in silico* prediction and perturbation. To train this highly parameterized model using densely sampled time-series data we developed a method for continuously collecting scRNA-seq data for budding yeast (*Saccharomyces cerevisiae*). We collected 173,000 scRNA-seq transcriptomes during response to treatment with the Tor Complex 1 (TORC1) inhibitor rapamycin, trained models to predict mRNA production and decay for each transcript, and learned the regulatory relationships between TFs and transcripts. We validated these estimates using scRNA-seq data collected at discrete timepoints, and prediction of gene expression states from *in silico* TF perturbation. Our method establishes a framework for efficient and accurate construction of GRNs, estimation of RNA kinetic parameters, and prediction of gene expression states using scRNA-seq data.

## 2 Results

Saccharomyces cerevisiae responds to the TORC1 inhibitor rapamycin through transcriptional regulation and altering regulated mRNA decay (*36*). Using bulk RNA-seq we confirmed that within five minutes of addition of rapamycin to exponentially-growing yeast in rich media, the abundance of stress response and metabolic transcripts increases and the abundance of ribosomal transcripts decreases (Figure 1A). The rate of change in transcript abundance during this response can be modeled for each gene using the ordinary differential equation (Equation 1).

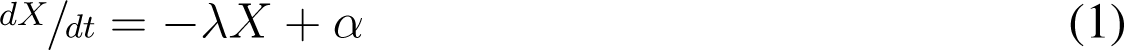

**Figure 1:**
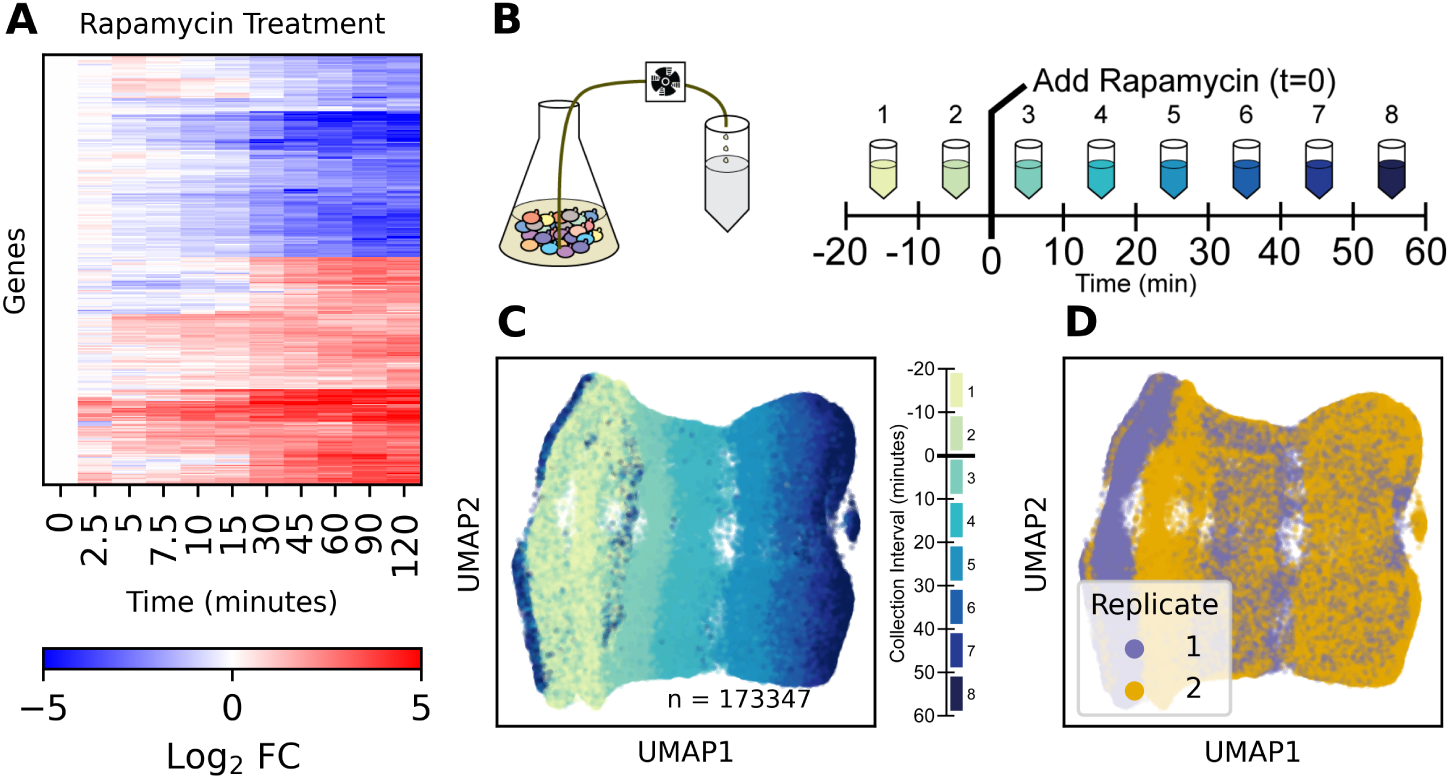
Time-dependent response to the TORC1 inhibitor rapamycin. (**A**) Bulk RNA-sequencing transcriptome response to rapamycin (Log_2_ Fold Change compared to untreated time 0; n=3). (**B**) Schematic of continuous-sampling single-cell RNA sequencing. Cells are collected for 10 minutes into mixing saturated ammonium sulfate and sequenced using the 10x genomics gene expression system. (**C**) 2D UMAP plot of individual *Saccharomyces cerevisiae* cells responding to rapamycin treatment, colored by sampling time. (**D**) 2D UMAP plot as in **C** colored by experimental replicate.

Here, the rate of change ^dX^⁄dt of mRNA expression *X* with time *t* is a combination of a decay component *−λX*, where *λ* is the mRNA decay rate, and a production component *α*, where *α* is the rate of transcript production. The rate of mRNA decay and transcription vary over time and between transcripts. Transcription rates are regulated by transcription factors whereas mRNA decay rates are regulated by RNA binding proteins and nucleases. We sought to simultaneously learn the kinetic parameters for every gene and identify the regulatory relationships between transcription factors and their targets using an interpretable deep learning framework. Training this highly parameterized model is data-intensive, requiring suitable training data with high temporal resolution.

### 2.1 Measuring Continuous Single Cell Response to Rapamycin

To minimize the interval between individual data points, we developed an experimental method for densely sampling time series data. Prior to, and following treatment, we continually pumped culture into excess saturated ammonium sulfate to collect and fix cells (Figure 1B), collecting cells in separate samples over sequential 10 minute intervals. This sampling design captures the transcriptome of individual cells over a continuous temporal distribution, unlike a standard discrete time point sampling. Using scRNA-seq we quantified the transcriptome response to rapamycin in individual cells, assaying 173,361 cells in eight sequential 10-minute time intervals, in two biological replicate experiments (Figure 1C-D).

Summary Uniform Manifold Approximation and Projection (UMAP) plots of cell transcriptomes show that the strongest transcript expression differences between cells is due to rapamycin treatment and the cell cycle, with minimal impact due to batch effect or sequence count depth (Supplemental Figure 1). Cells respond to rapamycin by downregulating ribosomal transcripts and upregulating stress response transcripts (Supplemental Figure 1G-H) as expected. The gene expression response is highly reproducible between replicates, and is only observed in wild-type cells. *Fpr1*Δ cells, which are genetically resistant to rapamycin, do not exhibit a transcriptional response (Supplemental Figure 2).

### 2.2 Assigning Response Times to Cells

The rapamycin response and cell cycle programs have distinct temporal transcriptional profiles, which are separable by principal component analysis (Supplemental Figure 3). We grouped transcripts into response to rapamycin and cell cycle transcriptional programs based on cosine distance between transcript abundance (Figure 2A; Supplemental Figure 5-7). For each program, we calculated a low-dimensional time trajectory (Supplemental Figure 8-12, and assigned a response time to each cell (Figure 2B). Cell cycle time estimates are continuous until rapamycin is added; afterwards, some intervals in the cell cycle have few or no cells assigned (Figure 2B). We hypothesize that this is due to rapamycin treatment causing cells to accumulate at transcriptional states that correspond with cell cycle checkpoints.

**Figure 2:**
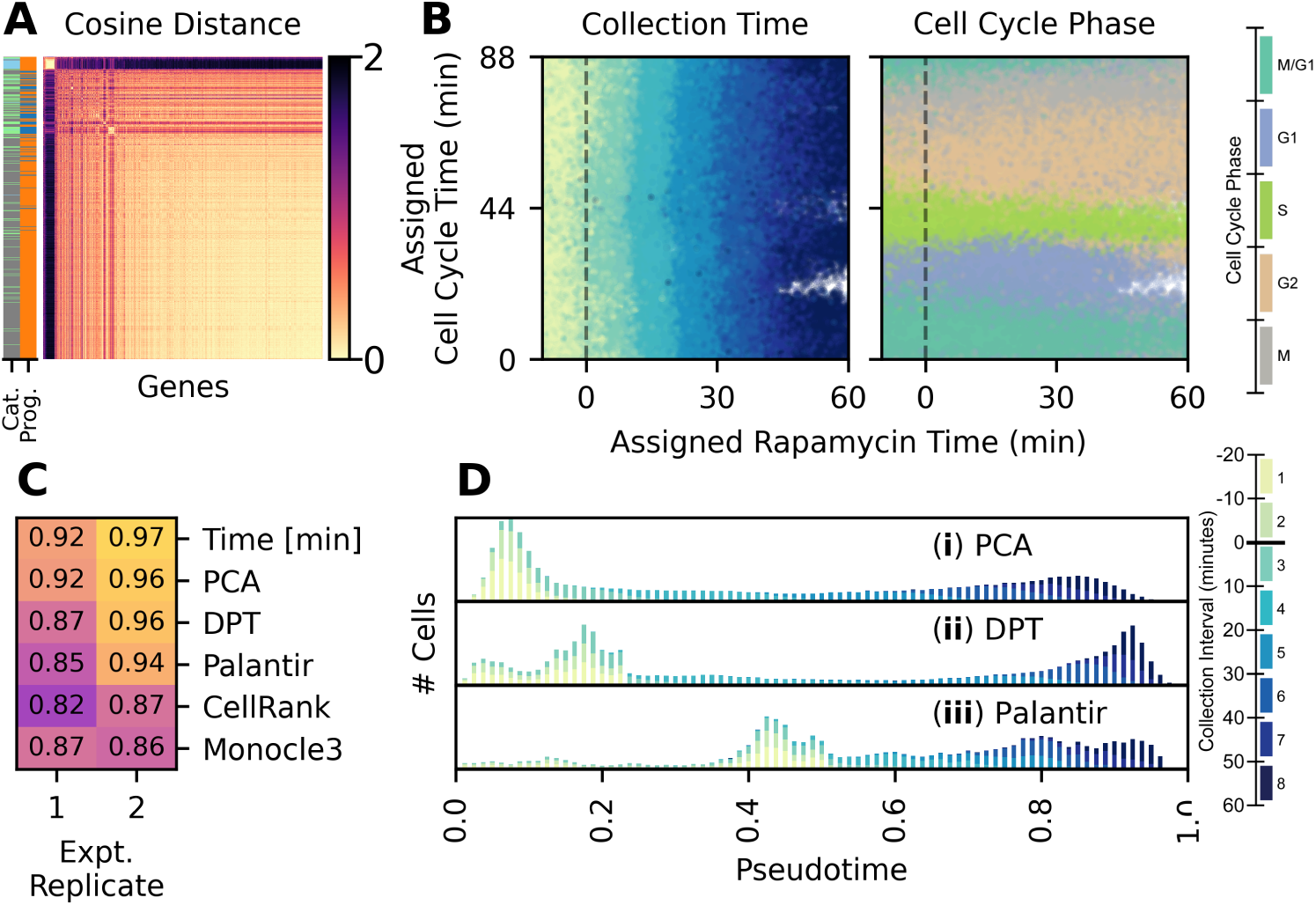
Assigning times to individual cells. (**A**) Cosine distance between transcripts. Transcripts are labeled by categories (Cat.), where green indicates cell cycle linked genes, blue indicates ribosomal genes, and gray is other genes. Transcripts are also labeled by program (Prog.), where orange is the response to rapamycin program and blue is the cell cycle program (**B**) Time values assigned to each wild-type cell. Cells are colored by collection interval (left) and cell cycle phase (right). (**C**) Performance of temporal and published pseudotemporal methods (DPT, Palantir, CellRank, and Monocle3) at ordering cells. Numeric values are spearman rho correlation between assigned time values and the collection interval. Spearman rho correlations for DPT, Palantir, CellRank, and Monocle3 are the maximum values from a grid search for *k*-NN parameters *k* and *N PCs* (Supplemental Figure 13) (**D**) Histogram of cells over pseudotime, calculated with (i) min-max scaling of the first principal component, (ii) DPT, and (iii) palantir. DPT and palantir use *k*-NN parameters that maximize spearman rho correlation with collection interval.

We compared our method of temporal assignment to commonly used methods for assigning pseudotime. We calculated the accuracy of temporal or pseudotemporal time assignments by comparing the ordering of time assignments to the known collection interval using spearman rho correlation (Figure 2C). We find that linear principal component-based methods perform well, as do diffusion pseudotime (DPT) (*37*) and palantir (*38*), although DPT and palantir assign a wider range of pseudotimes to cells prior to rapamycin treatment (Figure 2D). By contrast, CytoTrace (*39*) relies on assumptions that do not hold for this experiment, and Monocle (*40*) has strong dependence on a UMAP embedding, which is unpredictably sensitive to k-nearest neighbour (*k*-NN) embedding parameters (Supplemental Figure 13).

### 2.3 Predicting Transcriptomes with Dynamical Network Inference

To explicitly model transcription as a time-dependent process we extended our static neural network framework for GRN inference (*35*) using a recursive neural network (RNN) (Figure 3A). This model framework requires a prior knowledge network of TF to gene relationships (Supplemental Figure 14), which we applied as a constraint mask so that nodes of the first hidden layer can be interpreted as TFs. We quantified the influence of a TF on expression of each transcript by removing the TF node and calculating the coefficient of partial determination for each output transcript node. We then defined regulatory edges in a GRN by ranking every TF to gene edge based on the coefficient of partial determination. We tested the ability of the model to identify novel TF to gene regulatory relationships by holding out a subset of known interactions in the constraint mask and computing the Area Under the Precision-Recall curve (AUPR) based on recovery of the held-out known interactions. For all deep learning models, we used the Adam optimizer (*41*) and selected the learning rate (*γ* = 5 *∗* 10*^−^*^5^) and weight decay (*λ* = 10*^−^*^7^) to maximize AUPR scored against held-out network information and to maximize the total coefficient of determination R^2^ (Supplemental Figures 15-16).

We find that deep learning models greatly outperform linear models at recovering known regulatory TF to gene relationships (Figure 3B). Furthermore, the dynamical RNN models can make accurate forward predictions in time while remaining biologically interpretable, with only a modest decrease in GRN recovery performance (AUPR). To test the predictive power of the trained model, we used as input data the transcriptomes of untreated cells and iteratively predicted the full time course of gene expression in response to rapamycin treatment (Figure 3C).

**Figure 3:**
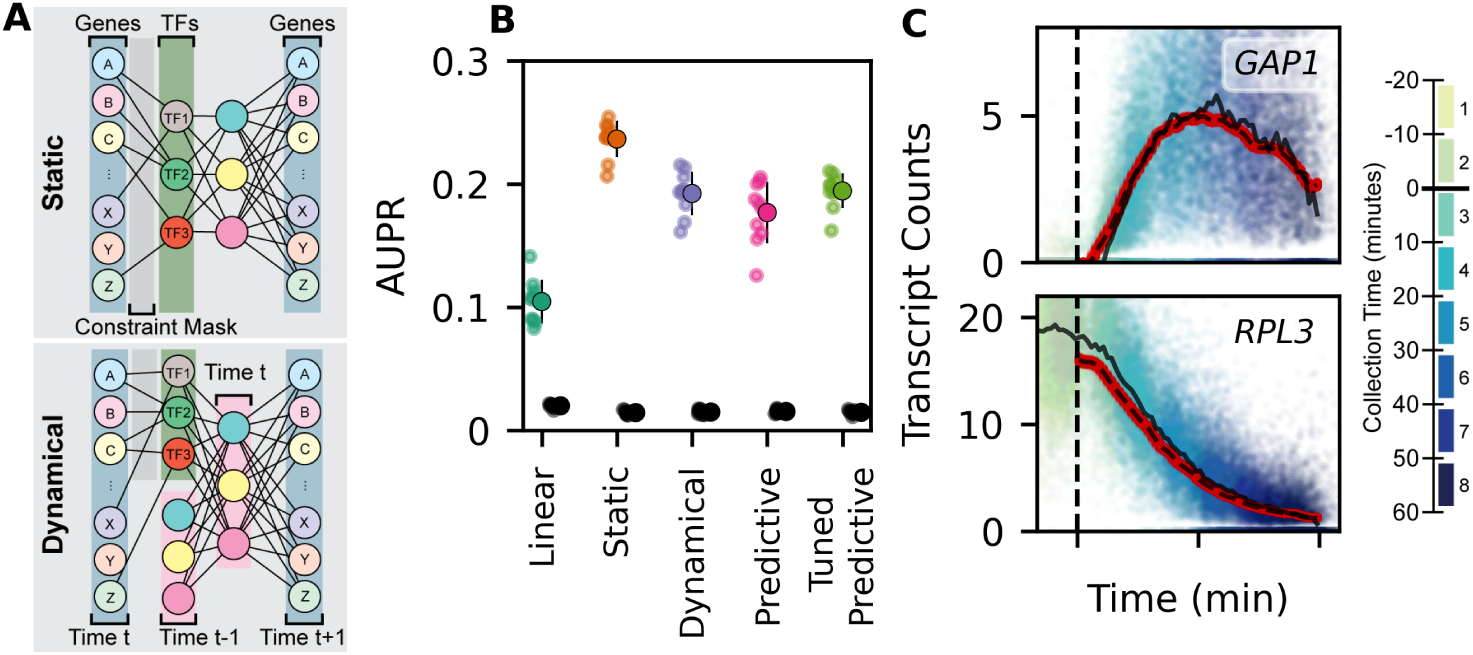
Dynamical Deep Learning Network Inference and Count Prediction. (**A**) Model schematics of static (upper) and dynamical (lower) deep learning model. Both models are interpretable for TF to transcript regulation as a consequence of the prior knowledge constraint mask. (**B**) Model performance computed as Area Under the Precision-Recall curve (AUPR), scored using transcripts that are held out of the prior knowledge constraint mask. Models scored are the Inferelator (*12*) with no time component (Linear) (*12*), deep learning with no time component (Static) (*35*), deep learning with a dynamic component (Dynamical), deep learning with a dynamic component trained only for long-term predictions (Predictive), and the Dynamical model fine-tuned for long-term predictions after initial training (Tuned Predictive). Black points are negative controls, where the labels on the prior knowledge constraint mask have been randomly shuffled. (**C**) Expression of the upregulated amino acid permease *GAP1* and the downregulated ribosomal protein *RPL3* plotted against rapamycin response time for each cell held out of the training data for validation (n=40633). Cells are colored by collection interval. Red points are predicted expression (from 0 to 60 minutes of rapamycin treatment), starting from untreated cells provided as input to the trained model (Tuned Predictive). Solid black lines are the medians of observed counts over time. Dashed black lines are the medians of predicted counts over time (red points).

### 2.4 Predicting RNA Velocity

Simultaneous estimation of the GRN structure and the RNA kinetic parameters corresponding to production and decay rates (Eq. 1) requires first estimating the RNA velocity, here defined as the rate of change in expression (dX/dt) of each transcript. We first made a preliminary estimate of RNA velocity by applying a self-supervised denoising method (*42*) and embedding cells into a *k*-NN graph. For every cell *i*, we defined a local neighborhood of connected cells, calculated the change in transcript abundance (dX) in the local neighborhood compared to cell *i*, and regressed it against the change in time (dt) to estimate RNA velocity (dX/dt) for each transcript (Supplemental Figure 17-18). We used the resulting RNA velocity estimate to train a new model that outputs RNA velocity and selected optimal hyperparameters based on cross-validation of held-out information (Supplemental Figures 19-20).

As with the RNN model that predicts count data, the first hidden layer corresponds to TFs, and the GRN is defined using coefficients of partial determination for each transcript. We find that models that predict RNA velocity recover known regulatory TF to gene relationships (Figure 4A) although GRN performance is slightly reduced relative to directly modeling count data. By predicting RNA velocity at time *t* from input transcript counts at time *t* (Figure 4B), and then updating transcript expression using RNA velocity, the model predicts the expected counts at *t* + 1 (Figure 4C), allowing for iterative forward prediction of RNA velocity. This predictive velocity model accurately reconstructs the overall trajectory of gene expression in response to rapamycin when untreated cells are provided as input (Figure 4D)

**Figure 4:**
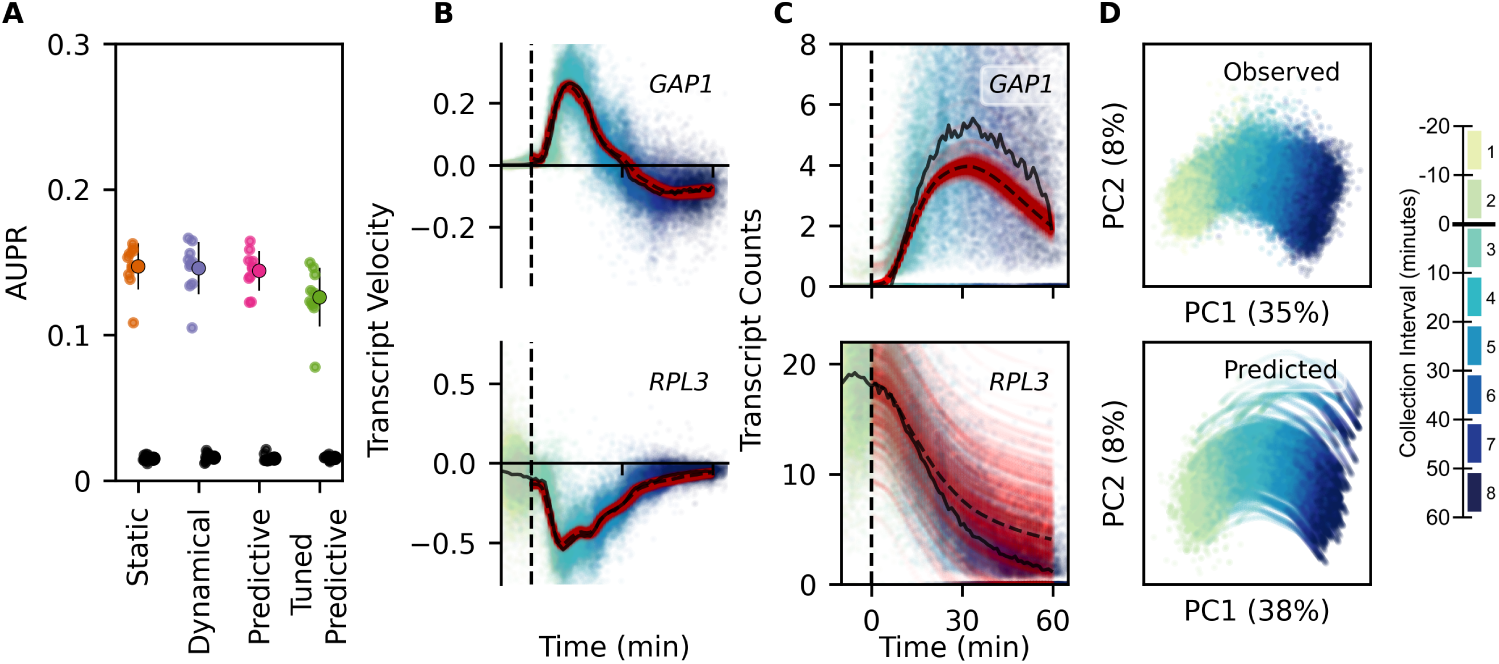
Dynamical Deep Learning Network Inference And Velocity Prediction. (**A**) Model performance (AUPR), scored using genes that are held out of the prior knowledge constraint mask. GRN is constructed by ranking on coefficient of partial determination to the primary model output, RNA velocity. (**B**) RNA velocity of *GAP1* and *RPL3* plotted against rapamycin response time and colored by collection interval for each cell held out of the training data for validation (n=40633). Red points are predicted velocity, starting from untreated validation data cell counts provided as input to trained model. Solid black line is the median RNA velocity of cells held out of training as a validation set. Dashed black line is the median predicted velocity prediction. (**C**) Expression of *GAP1* and *RPL3* plotted against rapamycin response time and colored by sample collection interval for each validation data cell. Predictions and medians over time are plotted as in **B**. (**D**) Principal component plot of observed cell transcriptomes held out of the training data for validation (n=40633), and cell transcriptomes (n=30170) from predicted trajectories (0 to 60 minutes) of rapamycin treatment.

### 2.5 Decomposing RNA Velocity into RNA Transcription Rates and Decay Rates

We further separated RNA velocity into a production and degradation component (Eq. 1) by extending our neural network model to include a decay module *f_λ_*(*t*) which predicts decay rates, and a transcription module *f_α_*(*t*) which predicts rates of transcription production (Figure 5A). Both model modules are trained to minimize velocity error, as well as an additional training constraint on the decay module to minimize distance from a lower-bound estimate of decay rate (Supplemental Figure 21-23). The first hidden layer of the transcription model is interpreted as transcription factors, allowing for construction of a transcriptional GRN.

**Figure 5:**
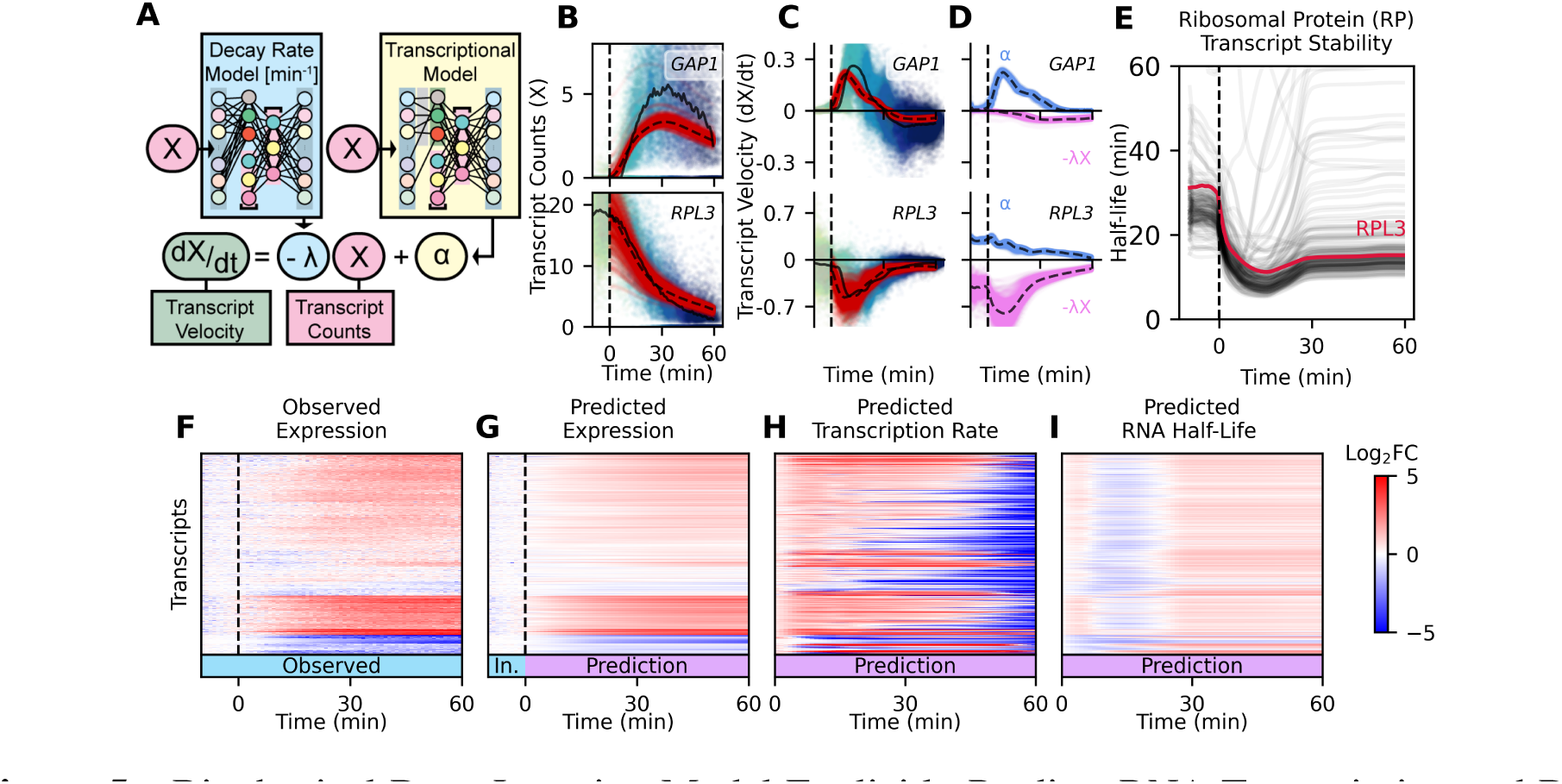
Biophysical Deep Learning Model Explicitly Predicts RNA Transcription and Decay. (**A**) Schematic of deep learning model that predicts transcriptional output and mRNA decay rates over time (**B**) Expression of *GAP1* and *RPL3* plotted against rapamycin response time and colored by collection interval for each validation data cell. Red points are predicted velocity, starting from untreated validation data cell counts provided as input to trained biophysical model. Solid black line is the median RNA velocity of cells held out of training as a validation set. Dashed black line is the median of the expression prediction. (**C**) Velocity of *GAP1* and *RPL3* plotted as in **B**. (**D**) Velocity predictions from **C** separated into positive transcriptional velocity *α* (blue) and negative decay velocity *λ*X (magenta) components. (**E**) Decay rate model predictions, converted to half-lives, for all 152 ribosomal protein (RP) transcripts. Plotted as the mean of all predicted decay rates for each transcript at each time. *RPL3* half-life highlighted in crimson. (**F-I**) Log_2_ fold change heatmap of observed transcript expression (**F**), predicted transcript expression (**G**), predicted transcription rate (**H**), and predicted mRNA half-life (**I**) during rapamycin treatment, compared to pre-treatment cells. Observed and model input data (In.) time intervals are labeled in blue, and model predictions are labeled in purple. An increase in RNA half-life (red) means RNA decays more slowly, and is more stable.

A biophysical RNN model predicts RNA velocity and uses those velocity predictions to predict transcript expression over time (Figure 5B-C). It further separates RNA velocity into a positive transcription rate component, and a negative decay component for each transcript (Figure 5D). The decay component of the RNA velocity module is the product of a predicted decay rate (*λ*) and the transcript expression (*X*). Using this model, we determined changes in mRNA stability over time and found that it is consistent with regulated changes in mRNA stability. For example, ribosomal protein (RP) transcripts are highly correlated during rapamycin response (Figure 5E). The model predicts that RP transcripts have a half-life of 20-30 minutes prior to TORC1 inhibition by rapamycin, but are subsequently destabilized with a half-life of 10 minutes. Predicted transcript expression, made by iteratively predicting 60 minutes of response to rapamycin treatment from untreated cells, matches the observed transcriptome well (Figure 5F-G), and both transcription rate and decay rates are temporally regulated (Figure 5H-I).

### 2.6 *In silico* Perturbation of RNA Transcription Model

The use of a neural network structure that reflects a GRN structure, in which nodes in the hidden layer are assigned a TF identity, enables additional analyses. First, the value of each node is constrained to be positive with an activation function, and these values are interpretable as Transcription Factor Activity (TFA). TFA quantifies the influence that a TF has on the transcription rate of its targets, accounting for the multiple ways in which the TFs are regulated, such as expression, post-translational modifications, nuclear localization, and cofactor binding. We found that the model identified core ribosomal TFs (*RAP1*, *SFP1*) and glycolytic TFs (*GCR1*) as highly active in proliferating cells (Figure 6A). When TORC1 is inhibited by rapamycin, these TFs are predicted to have sharply decreased activity, and other TFs are activated, such as those responsible for stress response (*GCN4*) and nitrogen metabolism (*GLN3*).

**Figure 6:**
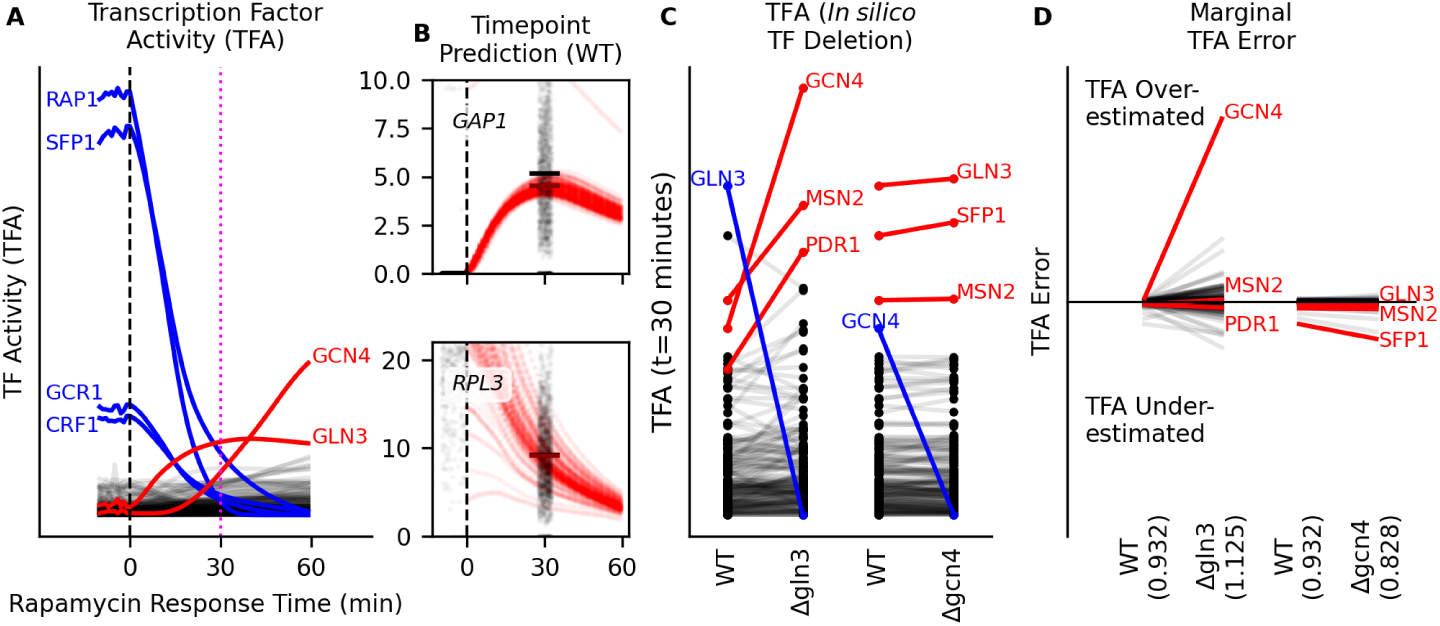
TF activity and *in silico* TF Perturbation Modeling. (**A**) TF activity predictions for 60 minutes of rapamycin treatment, starting with untreated, wild-type cells (*43*) (GSE125162) as input to the biophysical model (Figure 5). A rapamycin response timepoint for comparison was collected at 30 minutes (magenta dashed line). (**B**) Comparison between predicted response to rapamycin treatment (red points) and observed response to rapamycin treatment (black points, jittered for display), measured at t=30 minutes. (**C**) Comparison of predicted TF activities 30 minutes after rapamycin treatment between wild-type (WT) and *in silico* deletions (gln3Δ, gcn4Δ). Blue TFs have their activity set to zero at all times. (**D**) Predicted expression from wild-type and *in silico* gln3Δ and gcn4Δ modeling are compared to observed expression after 30 minutes of rapamycin treatment. Value in parentheses on the x-axis is the loss (mean squared error) of model predictions to observed values. TFA marginal model error is defined as the partial derivative of loss with respect to TFA, multiplied by TFA. Marginal model error is summed over time and then averaged over all observations.

Second, our modeling approach means that TF activity can also be altered *in silico* to predict the consequences of the alteration. We tested the utility of *in silico* predictions by fixing a TF’s activity to a specific value in the TF activity layer (i.e. 0 for a TF deletion) and then predicting the transcriptome response to rapamycin over time in the absence of the TF. We evaluated *in silico* predictions using an independent data set (*43*) that contains 12 different genotypes (wild-type and 11 TF deletions) that were assayed using scRNA-seq following 30 minutes of rapamycin treatment. We provided the model with data from untreated cells as input, and compared the predictions at 30 minutes following rapamycin addition to the observed values (Figure 6B).

In the absence of *GLN3* the transcriptional response to rapamycin treatment accurately predicts a reduction in expression of some of its known targets (gln3Δ; Supplemental Figure 24). Loss of *GLN3* is also predicted to lead to a compensatory increase in the activity of the stress-response TFs *GCN4*, *MSN2*, and *PDR1* (Figure 6C). The predicted increased in activity of these TFs to compensate for *GLN3* deletion is likely due to *GLN3* having many TF-TF interactions (e.g. protein-protein interactions or competitive binding). By contrast, the loss of *GCN4* has little effect on other TF activities, which we interpret as evidence of few *GCN4* TF-TF interactions when responding to rapamycin treatment.

We evaluated *in silico* TF deletions by comparing predicted counts to observed counts when the TF is deleted (t=30 minutes). The gln3Δ predictions (1.13) are less accurate overall than the wild-type predictions (0.93) using mean squared error, but gcn4Δ predictions are more accurate (0.828). This overall model error is backpropagated to the TF hidden layer to get marginal error for each TF. The increase in *MSN2* and *PDR1* activity when *GLN3* is deleted is real, as there is low marginal error for those TFs when comparing to observed gln3Δ cell expression, but there is high marginal error for *GCN4*, which indicates that the predicted increase in *GCN4* activity is incorrect (Figure 6D).

## 3 Discussion

To date, numerous GRN inference methods have been developed for identifying the transcriptional regulators of specific genes. However, predicting the consequences of genetic or environmental changes on gene expression, and estimation of the changes in rates of production and degradation that mediate those changes, presents a more challenging problem that has remained unsolved. Biophysical models that can be used for *in silico* experimentation are a necessary step to extend GRNs beyond descriptive models of gene expression.

One recent approach has been to simplify this task by predicting changes in a low-dimensional embedding (*13*) which represents cell developmental state. Although this is a powerful approach, predictions are not interpretable as biophysical processes, and the molecular mechanisms that underlie the changes to low-dimensional trajectories must be determined experimentally. To address this limitation, we have developed a GRN model that makes explicit predictions about mRNA transcription and decay rates for the entire transcriptome and defines transcriptional regulatory relationships. The modeling framework facilitates *in silico* prediction, perturbation, and analysis.

A key feature of our modeling framework is the quantification of expression dynamics as RNA velocity and its decomposition into a production and degradation component. This is possible due to the extremely high temporal resolution afforded by our novel experimental method for collecting transcriptomes at high resolution. However, our modeling framework is equally applicable to data collected using standard time course experiments, with lower temporal resolution. Estimation of transcriptional output and mRNA decay rates over time make clear that both of these kinetic parameters are altered to enact transcriptome remodeling, and a complete model of gene expression must incorporate both.

We constrain the transcription model based on existing knowledge, which facilitates interpretation of the first hidden layer as TFs that control transcript production. At this point, regulation of mRNA decay is not assigned to specific regulatory proteins, as we lack sufficient existing knowledge about the decay regulatory network to constrain the decay model. This should be addressable in the future through incorporation of additional experimental evidence enabling causal predictions about decay regulation, and *in silico* perturbation of decay factors.

The most interesting application of a deep learning GRN framework is building high-quality models from training data collected for the express purpose of optimizing modeling, and applying those models to make predictions from other samples. Here we show that this strategy can be used to train a model that predicts response to rapamycin and then apply that model to non-dynamic observational data (*43*) to construct response trajectories. We envision fine-tuning existing models, instead of building new models for every project, and can foresee using pooled single-cell perturbation experiments (*43–45*) to create a pre-trained regulatory model that is the base for future regulatory inference. These are major modeling advantages, and our expectation is that deep learning models will replace traditional methods of GRN inference in the future.

## 4 Methods and Materials

All materials are released under CC-BY 4.0 and all code is available under the permissive MIT or BSD licenses. Single-cell RNA-sequencing (scRNA-seq) data has been deposited in NCBI GEO under the accession number GSE242556. Code to assign cells to times and to calculate RNA velocity is available on GitHub and PyPi (inferelator-velocity v1.0.0). Supirfactor-dynamical code for modeling is available on GitHub and PyPi (supirfactor-dynamical v1.0.0). All figures are generated with matplotlib (*46*) in python and code is available on GitHub.

### 4.1 Yeast Strain Construction and Growth

All yeast strains used in this work are derived from the prototrophic FY4 (MATa) and FY5 (MATα) parental strains (*47*). Strains are created by transformation using the standard lithum acetate transformation protocol (*48*). Yeast strains and oligonucleotide sequences are provided in Supplemental Tables. Unless otherwise noted, all strains are grown in YPD (2% w/v Bacto Peptone, BD Biosciences #211677; 2% w/v D-glucose, BD Biosciences #215520; 1% w/v Yeast Extract, BD Biosciences #212750) at 30°C.

#### 4.1.1 Construction of Barcoded Strains

Barcodes are located in the 3’ Untranslated Region (UTR) of the geneticin (G418) resistance cassette and the 3’ UTR of the nourseothricin (NAT) resistance cassette. A library of plasmids containing unique barcode sequences was generated (*43*) and used as the template for PCR with primers containing gene-specific targeting homology arms (1x Q5 Master Mix, New England Biolabs #M0494S; 1 ng template plasmid; 250 nM/each primer). The PCR-amplified homology-directed repair cassette was then transformed into yeast. The G418-resistance cassettes were transformed into FY4 (MATa) and selected on YPD + G418 (400 µg/mL; Gold Biotechnology #G-418-5) plates. The NAT-resistance cassettes were transformed into FY5 (MATα) and selected on YPD + NAT (200 µg/mL; Gold Biotechnology #N-500-1) plates. Correct site integration was validated by colony PCR using primers specific for the integration site and for the resistance cassette. Colony PCR products were cleaned using a silica spin column (QIAGEN #28104; manufacturer’s protocol) and sanger sequenced (GeneWiz) to determine the sequence of the strain-specific molecular barcode.

An array of 72 uniquely-identifiable wild-type strains was generated by crossing 6 FY4 strains where the inactivated mating-type switching loci *HO* was replaced by a uniquely-barcoded G418-resistance cassette and 12 FY5 strains where the *HO* loci was replaced by a uniquely-barcoded NAT-resistance cassette. MATa and MATα *HO*.6.-barcoded strains were arrayed such that each well contained a unique combination of G418 UTR barcode and a NAT UTR barcode. MATa and MATα strains were separately grown overnight in YPD in a 96-well plate (Corning #3788; 150 µL media / well), sealed with a gas-permeable membrane (Diversified Biotech #BERM-2000). For each well, 25 µL of MATa culture was added to 25 µL of MATα culture in 100 µL fresh YPD and grown for 6 hours at 30°C. The 96-well plate was centrifuged to pellet cells and the pellets were pinned to a YPD + G418 + NAT plate (Nunc Omnitray #165218) and grown 2 days at 30°C to select for mated MATa/α diploids. These diploids were pinned to a second YPD + G418 + NAT plate for a second round of diploid selection and then pinned into YPD in a round-bottom plate (Corning #3788; 150 µL media / well), sealed with a gas-permeable membrane (Diversified Biotech #BERM-2000), and grown with shaking at 30°C overnight. Cells were pelleted and the supernatant media was aspirated and replaced with 150 µL of 50% glycerol (Ricca Chemical # R3290000) for long-term storage in a -80°C freezer.

The *fpr1*.6. mutant cells were created using the same transformant protocol as above. Six MATa *fpr1*.6. strains with barcoded G418-resistance cassettes were crossed to an MATα *fpr1*.6. strain using a NAT-resistance cassette with the mating protocol above. They were selected as above and stored in a -80°C freezer.

#### 4.1.2 Growth Rate

Growth of these strains was measured by inoculating 50 mL of YPD with 10^8^ cells from overnight culture, growing for 3 hours, and then taking cell density measurements (cells/mL) with a coulter counter (Beckman Coulter Z2 Particle Counter #6605700) every 10 minutes. Doubling time was calculated by regressing log_2_(cell density) against time.

#### 4.1.3 Bulk RNA-sequencing Culture & Sample Preparation

Three wild-type FY4/5 (MATa/MATα) replicates were inoculated into YPD and grown overnight at 30°C. Overnight cultures were subcultured 1:100 into 25 mL YPD and grown 4 hours at 30°C with shaking. A 0 minute timepoint was taken by flash spinning 1 mL culture in a microfuge tube, discarding media, and snap-freezing in liquid nitrogen. 200 ng/mL rapamycin was added to the culture (Millipore #553210; stock solution 1 mg/mL in ethanol) and culture was collected and snap-frozen as before at 2.5, 5, 7.5, 10, 15, 30, 45, 60, 90, and 120 minute timepoints of rapamycin treatment. Cell pellets were stored at -80°C until extraction.

#### 4.1.4 Culturing and Continuous Sampling

Frozen (−80°C) glycerol stocks were pinned onto a YPD + NAT plate and grown at 30°C. The wildtype (*ho*.6.) strains and the *fpr1*.6. strains from separate freezer plates were spotted into separate positions onto a single YPD + NAT plate from frozen stocks. These strains were then pinned into YPD + G418 in a 96-well plate (150 µL media / well), sealed with a gas-permeable membrane, and grown with shaking at 30°C overnight. Overnight cultures were pooled from a 96-well plate into a single 50 mL conical, washed twice with 25 mL YPD, and resuspended in 10 mL of YPD. 10^9^ cells were used to inoculate a 1L baffled flask containing 500 mL prewarmed YPD which was grown with shaking for 4 hours at 30°C.

At 4 hours, a stir bar was added to the flask, and it was transferred onto a stir plate in a 30°C environmental chamber. A weighted sinker attached to 1/8” tubing was dropped into the flask and attached to a peristaltic pump (Watson-Marlow Cartridge Pump, 32 Channel, 205S / CA32) running at 60 RPM (0.6 mL / min) for sampling. Samples were collected continuously into 50 mL conicals containing 20 mL RNAlater (saturated ammonium sulfate, 20mM EDTA, pH 5.2; QIAGEN #76106) and a stirring magnetic stir disc (V&P Scientific #772DP-N42-5-2). Sample collection conicals were changed every 10 minutes and the collected samples were immediately placed on ice. After 20 minutes, 200 ng/mL rapamycin was added to the stirring culture (Millipore #553210; stock solution 1 mg/mL in ethanol). The collection tubing was disconnected from the sampling conical, 8 mL of volume was drawn through using a syringe (2^°^x the 4 mL dead volume of the sampling tubing), and the sampling tubing was immediately reconnected for continued collection. Sampling continued for 60 additional minutes, yielding 2 10-minute sample pools without rapamycin treatment and 6 10-minute sample pools with rapamycin treatment. Cells were pelleted, washed once with 1 mL RNAlater and transferred to a microfuge tube, and then resuspended in 500 µL RNAlater. Samples were stored at -20°C after determining their density with a coulter counter.

### 4.2 Sequencing Sample & Library Preparation

#### 4.2.1 Bulk RNA-sequencing Sample & Library Preparation

RNA was extracted from from frozen cell pellets by adding 0.5 mL TRIZOL (ThermoFisher #15596026) and 100 µL chloroform. The aqueous layer was extracted again with 500 µL acid phenol chloroform (ThermoFisher #AM9720; pH 4.5), and that aqueous layer was extracted again with 500 µL chloroform. RNA was precipitated with 2.5x volumes ice-cold ethanol and 0.1x volume 3M sodium acetate pH 5.5 (ThermoFisher #AM9740) overnight at -80°C. RNA pellets were washed once with 750 µL ice-cold 70% ethanol and resuspended in 100 µL EB (10 mM TRIS pH 8, 0.05% TWEEN-20). RNA was quantified by Qbit and diluted to 10 ng/µL.

RNA was primed with dT30VN oligonucleotides that contain unique molecular identifiers (UMIs), sample-specific barcodes to facilitate pooling, and a common PCR handle sequence. Reverse transcription was performed by mixing 10 µL RNA (100ng) with 10 µL RT mix (5 µL 5x Maxima RT Buffer, 1.25 µL 10 mM/each dNTP [New England Biolabs #N0447S], 0.5 µL Lucigen NxGen RNase Inhibitor [Lucigen #30281–1], 0.5 µL 50 µM Template Switch Oligo [IDT], 0.5 µL 50 µM Barcode/UMI/poly-dTVN Oligo [IDT], 0.5 µL Maxima H Minus Reverse Transcriptase [ThermoFisher #EP0752], 6.75 µL H_2_O) and incubating at 53°C for 1 hour. Reactions were then heat-inactivated for 5 minutes at 85°C.

12 cDNA reactions were pooled so that each sample-specific barcode was unique in each pool, and the cDNA was purified with 1x volume of AMPure XP beads (Beckman Coulter #A63880) into 40 µL EB. 60 µL whole-transcriptome amplification (WTA) master mix (50 µL 2x KAPA HiFi Hotstart Readymix [Roche #KK2602], 1 µL 100 µM Forward Oligo, 1 µL 100 µM Reverse Oligo, 8 µL H_2_O) was added to each reaction and amplified using 8 cycles of PCR using 10 cycles of PCR (98°C for 3:00; 12 cycles of 98°C for 0:20, 55°C for 0:20, and 72°C for 1:15 min; 72°C for 3:00 min). WTA reactions were purified with 0.6x volume of AMPure XP beads, eluted into 25 µL EB, and quantified using a high sensitivity D5000 ScreenTape (Agilent #5067–5592).

Sequencing libraries were prepared by tagmenting 10 ng WTA DNA with Nextera XT kit (Illumina #FC-131-1024). Indexing PCR was primed with a nextera i7 index primer (e.g. N701-712) and a custom truseq i5 index primer specific for the PCR handle added onto the 3’ end of transcripts during reverse transcription, and amplified for 10 cycles. Sequencing libraries were verified using an Agilent 4200 TapeStation and D1000 tape (Agilent #5067-5582) and quantified by qPCR using a Roche LightCycler 480 and Illumina Library Quantification Kit (Kapa Biosystems #KK4854) Libraries were pooled evenly sequenced per NovaSeq XP flow cell (Illumina #20043130) using a 28bp read 1, 91bp read 2, 8bp index 1, 8bp index 2 run configuration.

#### 4.2.2 Single-Cell RNA-sequencing Sample & Library Preparation

5 * 10^6^ cells were aliquoted from the -20°C RNAlater samples. These cells were pelleted and washed once with spheroplast digest buffer (50 mM Sodium Phosphate pH 7.4, 1M Sorbitol, 1TT, 100 µg/mL BSA). After washing, cells were resuspended in 100 µL spheroplast digest buffer with 1U/mL Zymolyase T100 (Zymo Research #E1005) and incubated at 37°C for 20 minutes. Pools 4-8 from experimental replicate #2 were resuspended in 100 µL spheroplast digest buffer with 2.5U/mL Zymolyase T100 (Zymo Research #E1005) and incubated at 37°C for 20 minutes. After digest, 400 µL of RNAlater and 2 µL of 20 mg/mL BSA (New England Biolabs # B9000S) were added to the samples and incubated on ice for 5 minutes. The supernatant was carefully aspirated after pelleting cells at 375xg for 3 minutes at 4°C. Cells were then washed twice with 750 µL and then resuspended in 100 µL of spheroplast wash buffer (10 mM TRIS pH 8, 1M Sorbitol, 400 µg/mL BSA). Concentration of spheroplasted cells was determined by counting with a Fuchs-Rosenthal hemocytometer (InCyto #DHC-F01), and concentration was adjusted to 4 * 10^6^ cells/mL by adding additional buffer.

Single-cell gene expression was performed using a Chromium Next GEM Single Cell 3’ Kit v3.1 (10x Genomics #1000268). 69 µL of Reverse Transcription Master mix was prepared per-sample using the 10x kit reagents (18.8 µL RT Reagent B, 8.7 µL RT Reagent C, 2.4 µL Template Switch Oligo, 2 µL Reducing Reagent B, 37.2 µL Nuclease-free water). To this, 6 µL of 4 * 10^6^ cells/mL (24,000 total cells) was added, loaded into a single-cell microfluidic chip (10x Genomics #1000127) and run on the 10x Genomics Chromium Controller. The resulting single-cell emulsion was processed using the manufacturer’s protocol, with 9 cycles of PCR whole transcriptome amplification. Whole transcriptome amplifications were verified using a 4200 TapeStation and high-sensitivity D5000 tape (Agilent #5067-5592). 25% of the amplified whole transcriptome was used to generate sequencing libraries according to the 10x genomics library preparation kit protocol. In short, the samples were fragmented, adaptor ligated, and indexed with 10 cycles of PCR to add unique dual i5/i7 indexes (10x Genomics #1000215).

Strain-specific barcodes were extracted by PCR using the amplified whole transcriptome as template (1x KAPA HiFi Hotstart Readymix, Kapa Biosystems #KK2602; 200 nM/each primer; 1 µL10x whole transcriptome) with the following protocol: 98°C for 3:00; 6 cycles of 98°C for 0:20, 63°C for 0:20, and 72°C for 0:20 min; 72°C for 1:00 min. Separate PCR reactions were performed to extract the barcodes in the NAT^R^ and KAN^R^ UTRs. These reactions were cleaned up with 1x SPRIselect (Beckman Coulter #B23318), washed twice with 80% ethanol, and eluted into 20 µLof 10mM TRIS. This entire volume was then used as template for an additional i5/i7 indexing PCR reaction (1x KAPA HiFi Hotstart Readymix, Kapa Biosystems #KK2602; 200 nM/each primer; 20 µLbarcode amplicon elution) to add unique dual i5/i7 indexes (10x Genomics #1000215) using the following PCR protocol: 98°C for 0:45; 10 cycles of 98°C for 0:20, 54°C for 0:30, and 72°C for 0:20; 72°C for 1:00. Transcriptome and barcode amplicon library construction was verified using an Agilent 4200 TapeStation and D1000 tape (Agilent #5067-5582) and quantified by qPCR using a Roche LightCycler 480 and Illumina Library Quantification Kit (Kapa Biosystems #KK4854). Libraries were pooled to target 400 million reads per whole transcriptome library and 5 million reads per barcode amplicon library, allowing 4 lanes of 10x to be sequenced per NovaSeq S1 flow cell (Illumina #20028319) using a 28bp read 1, 91bp read 2, 10bp index 1, 10bp index 2 run configuration.

### 4.3 Sequencing Quantification

#### 4.3.1 Single-Cell RNA-sequencing Expression Quantification

Raw reads were demultiplexed from BCL into FASTQ files using bcl2fastq (Illumina) with run-appropriate settings and indexing run sheet. FASTQ files for each experimental sample were pseudoaligned and processed into a BUS file with kallisto bus (*49*), using an appropriate yeast reference transcriptome containing UTR sequences from earlier work (*43*), and appropriate chemistry configuration for a 10x v3.1 3’ gene expression library. BUS files were further processed with kallisto correct, in order to correct cell barcode reads to match the 10x genomics barcode whitelist provided with the CellRanger software package. This corrected BUS file was sorted with kallisto sort and counted with kallisto count.

Extracted barcode amplicon libraries were processed with a custom python script using the previously-published python fastqToMat0 package (*43*). In short, this package extracts cell barcodes and links them to transcriptional barcodes that identify the strain of the captured cell, while removing any cell barcodes which have multiple transcriptional barcodes (indicating that the cell barcode is a ‘doublet’ containing more than one cell). It then reads in the expression data generated by kallisto and produces a TSV file that includes strain metadata. This custom pipeline depends on the standard python numpy (*50*) and pandas (*51*) libraries. Additional experimental metadata was added to TSV files, and all samples were row-combined into a TSV file containing all expression data for this experiment.

Raw count data is denoted here as **Z** *∈* Z*^n×g,^*^+^ where *n* is cells (observations) and *g* is transcripts (features). This raw integer count data is standardized to **X** *∈* R*^n×g,^*^+^ by dividing all features of every cell by a cell-specific scaling factor, such that the sum of all counts in each observation is the same. These scaling factors were chosen so that all cells sum to the median number of counts per cell from **Z** (3099 counts/cell).

Published single cell data was obtained from NCBI GEO (GSE125162) with sra-tools’ prefetch and converted from SRA format to FASTQ format with fastq-dump. It was then processed as above into a standardized count matrix where all cells sum to 3099 counts/cell.

#### 4.3.2 Bulk RNA-sequencing Expression Quantification & Analysis

Bulk RNA-seq libraries were 3’ end sequenced with UMIs and sample-specific barcodes in Read 1 and transcript sequence in Read 2. They were therefore processed using the same pipeline tools as the single-cell expression quantification above. Each library was processed with bcl2fastq, kallisto, and bustools to count transcripts by UMI, yielding raw integer count data.

Bulk RNA-seq differential expression was performed with DESeq2 (*52*), testing for a fold change difference greater than 0.25x (log_2_ fold change threshold of 0.322), and rejecting the null hypothesis where the adjusted p-value was less than 0.01.

### 4.4 Gene Assignment to Programs

#### 4.4.1 Distance Metrics

For this analysis, **X** was log(x+1) transformed, and 2189 high-variance genes were retained, as defined by a minimum variance of 0.01 and minimum mean value of 0.0125 (using scanpy). Transformed expression data **X** was projected into 13 PCs, and then projected back into gene expression space, removing variance not captured in the first 13 PCs to give **X**_13PC_. 13 PCs were selected by molecular cross-validation (MCV) on raw UMI counts (*53*).

Distance was computed between genes *i* and *j* with corresponding expression vector **x***_i_* and **x***_j_* from **X**_13PC_, for euclidean distance (Eq. 2), cosine distance (Eq. 3), and manhattan distance (Eq. 4), and informational distance (Eq. 5) (*54*) (calculated with mutual information (Eq. 6) and joint entropy (Eq. 7)).

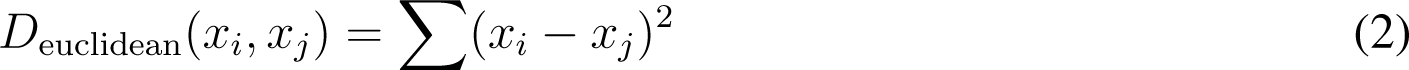

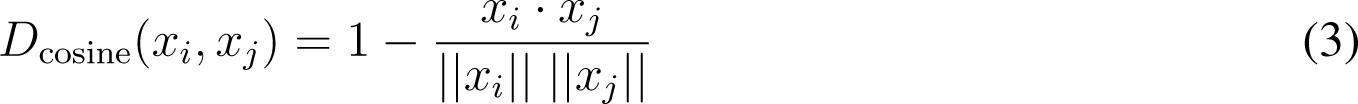

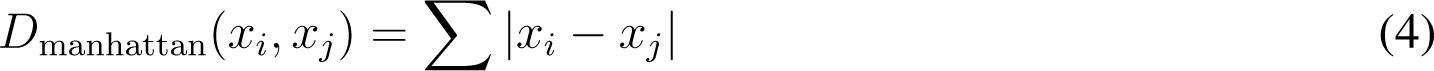

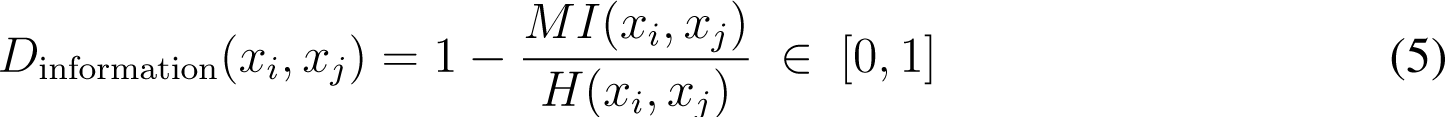

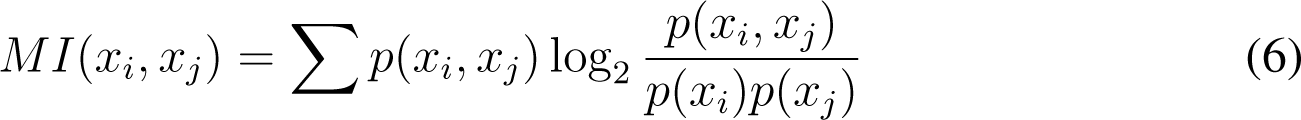

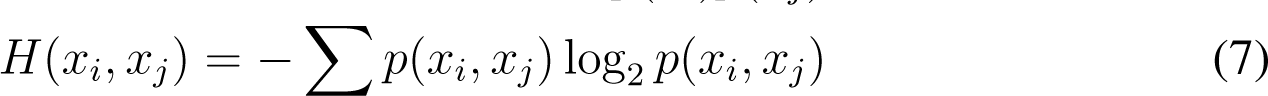

#### 4.4.2 Gene Clustering and Merging into Programs

We construct a gene-gene *k*-NN graph from **X**_13PC_ using cosine distance. Genes are clustered on the *k*-NN graph, with *k* = 21 with the leiden algorithm (*55*) (resolution of 1) into 15 clusters, each containing between 37 and 266 genes.

For each gene cluster *C ∈ {*1, 2*, …,* 15*}*, count data for only these genes, **Z***_C_* is selected, standardized, and log transformed independently. The highest explained-variance principal component (PC_1_) is calculated and used to ordering cells in cluster *C* on an equally spaced on a [0, 2*π*] interval. The *ρ*_circ_ correlation (*56*) between each gene cluster is then calculated (Eq. 8), and the distance between the gene clusters is defined as *D*_circ_ = 1 *− |ρ*_circ_*|*. These gene clusters are then further aggregated into two programs by agglomerative clustering using this distance metric, yielding 2038 genes in a rapamycin response program and 346 genes in a cell cycle program.

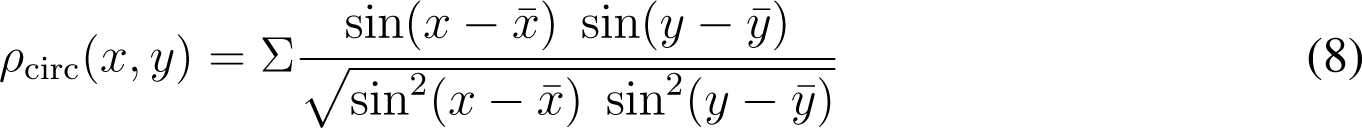

This analysis pipeline has been released as the programs module of the inferelator-velocity package.

### 4.5 Time Assignment to Cells

Times were assigned to wild-type and fpr1.6. cells separately, and to experimental replicates 1 and 2 separately, as follows. For each program, raw integer count data for the program genes, **Z**_P_, was standardized to the median count depth and then log(x+1) transformed into **X^°^**_P_. From **X^°^**_P_ we extracted 5-8 PCs (*N*), selected by MCV using **Z**_P_ as input, into **U**_N_.

For the rapamycin response program, each cell was assigned to an experiment sampling interval with known real-world times and order. Pools 1 & 2 are combined and assigned a centroid time of -5 minutes, pool 3 is 5 minutes, pool 4 is 15 minutes, pool 5 is 25 minutes, pool 6 is 35 minutes, pool 7 is 45 minutes, and pool 8 is 55 minutes. The centroid for each group of cells was identified in the PC space. From **U**_N_ we computed a cell-cell *k*-NN graph, *k* = 10, using euclidean distance. The distance between centroids of sequential groups was measured on the *k*-NN graph using Dijkstra’s algorithm (*57*).

To annotate each cell with a global time we applied the following procedure. For two connected groups *A* and *B*, all cells closest to the path between centroids *a* and *b*, *L_ab_*, are assigned a time *T_ab_*by rooting **U**_N_ in *a* and then projecting the resulting vector for each cell *u_c_*onto the vector *v_ab_*connecting *a* to *b*. The cell time, *T_ab_* is then the fractional distance between centroid times *t_a_* and *t_b_* formed by the projection, plus *t_a_* (Eq. 9), with *\.\* indicating the euclidean norm..

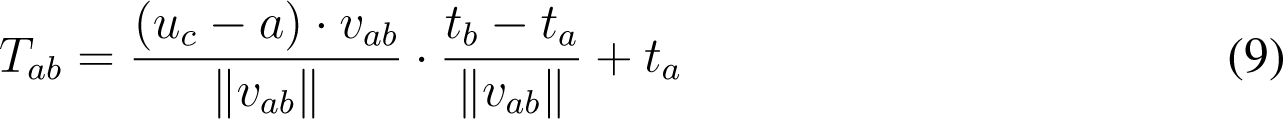

As an example, the centroids of pools 3 and 4 are connected by a 10 minute time vector, with a 5 minute offset from the first centroid. A cell that projects halfway (e.g. at 5 minutes) on the time vector between pools 3 and 4 will therefore be assigned a time of 10 minutes. This is repeated for all connected pairs of centroids to build times for all cells.

For the cell cycle program, cells were assigned to cell cycle phases based on existing published data (*58*) using an established method (*43*). The length of the cell cycle was set to 88 minutes based on an experimentally determined doubling time. Phase M1-G was assigned a centroid time of 7 minutes, G1 was assigned a centroid time of 22.5 minutes, S was assigned a centroid time of 39.5 minutes, G2 was assigned a centroid time of 56.5 minutes, and M phase was assigned a centroid time of 77.5 minutes. These phases were connected in a circle, and times were explicitly wrapped at 88 minutes. Time assignments to cells were done as for rapamycin response (Eq. 9)

#### 4.5.1 Pseudotime Calculations

For each pseudotime calculation, expression data was processed separately for each experiment and for wild-type and fpr1.6. cells. Pseudotime Principal Components (PCs) or Diffusion Components (DCs) hyperparameters were selected from 5, 15, 25, 35, 45, 55, 65, 75, 85, 95, or 105 components. Pseudotime *k*-Nearest Neighbors (*k*-NN) hyperparameters were selected from 15, 25, 35, 45, 55, 65, 75, 85, 95, or 105 neighbors. Monocle3 and palantir require a starting cell for pseudotime calculations. The cell with the most negative PC_1_ value was chosen and provided as a starting cell (cell 9214 for Expt 1 / WT, cell 3815 for Expt 1 / fpr1.6., cell 82689 for Expt 2/ WT, and cell 70873 for Expt 2 / fpr1.6.).

#### 4.5.2 Principal Component Pseudotime

Raw integer count data **Z** was standardized to the median count depth (3099 counts/cell) and then log(x+1) transformed to **X^°^**. Principal components were then calculated after centering and scaling **X^°^**to unit variance. The first principal component (PC_1_) was used to calculate pseudotime by interval normalizing to values from 0 to 1 ((Eq. 10)).

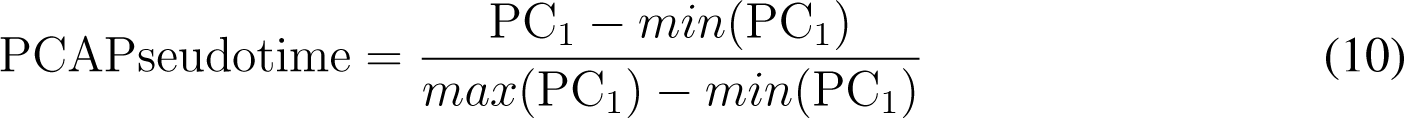

#### 4.5.3 Monocle3 Pseudotime

The R (v4.1.2) package Monocle3 (*19*) (v1.0.0; Commit# 004c096) was obtained from https://github.com/cole-trapnell-lab/monocle3. Raw integer count data **Z** was provided to the Monocle3 monolithic pipeline. In short, this pipeline normalizes, calculates *N* PCs, calculates a *k*-NN graph using *N* PCs, projects the cells into two dimensions with Uniform Manifold Approximation and Projection (UMAP) (*59*), clusters cells using the Leiden algorithm (*55*), and then assigns pseudotime values to each cell. Root cells were selected as noted above.

#### 4.5.4 Palantir Pseudotime

The python packages palantir (v1.0.0) (*38*) and harmonyTS (v0.1.4) (*60*) were obtained from the PyPi package repository with pip. Raw integer count data **Z** was standardized to the median count depth (with scanpy.pp.normalize per cell) and then log(x+1) transformed (with scanpy.pp.log1p). Palantir then calculates a diffusion map (using palantir.utils.run diffusion maps), modifies the diffusion map by force-directed layout (using harmony.plot.force directed layout), and calculates pseudotimes for each cell (using palantir.core.run palantir). Root cells were selected as noted above.

#### 4.5.5 CellRank (CytoTrace) Pseudotime

The python cellrank (v1.5.1) (*28*) package was obtained from the PyPi package repository with pip. Raw integer count data **Z** was standardized to the median count depth (with scanpy.pp.normalize per cell) and then log(x+1) transformed (with scanpy.pp.log1p). PCs (using scanpy.pp.pca) and a *k*-NN graph (using scanpy.pp.neighbors) were calculated, and data was imputed from scVelo (*29*) moments (using scvelo.pp.moments). Pseudotime was then calculated from a CytoTRACE (*39*) score (using cellrank.tl.kernels.CytoTRACEKernel).

### 4.6 Biophysical Transcriptional Expression Model

We fit transcriptional expression to a first order ordinary differential equation (Eq. 11) where *X ≥* 0 is transcript abundance in counts, *t* is time in minutes, Λ is RNA decay constant *λ_i_* for every gene *i* in minutes^-1^, and *α* is transcriptional production in counts *·* minutes^-1^.

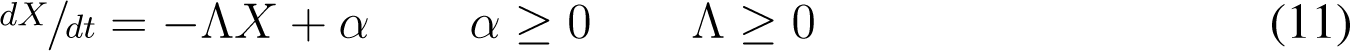

In this formulation, *X* is cells by genes, Λ is cells by genes, *α* is cells by genes, and decay and transcriptional production may vary from cell to cell.

#### 4.6.1 Denoising and Optimal *k*-NN Graph Construction

Denoising and *k*-NN graph construction is performed using the previously-published Denoising Expression data with a Weighted Affinity Kernel and Self-Supervision (DEWӒ KSS) package (*42*). This is based on a self-supervised method to construct a euclidean-distance weighted *k*-NN graph in a space defined by a specific number of PCs, where *k* is not constant and is allowed to vary per node. This method selects a *k*-NN graph that, when used for denoising, minimizes a mean squared error objective function based on noise2self (*61*). We use this as a principled way of constructing an optimal *k*-NN graph, without requiring *k* or the number of PCs to be set arbitrarily. Optimal *k*-NN graph construction and denoising was performed on wild-type and fpr1.6. cells separately, and on experimental replicates 1 and 2 separately.

#### 4.6.2 RNA Velocity Estimation

RNA velocity is the rate of transcript change over time. For each cell *c*, expression data of all cells connected to *c* in the optimal *k*-NN graph (**X***_c_*) were centered on the expression data of *c* to create a change in expression matrix Δ**X***_c_*. The time assigned to each connected cell *t_c_* is centered on the time of *c* to create a change in time matrix Δ*t_c_*. Δ**X_c_**is regressed against Δ*t_c_*(Eq. 12), and the estimated coefficients *β*^^^*_c_* (Eq. 13) are the rate of change of transcript over time and are interpreted as RNA velocity for cell *i*.

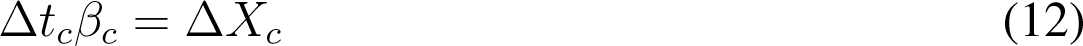

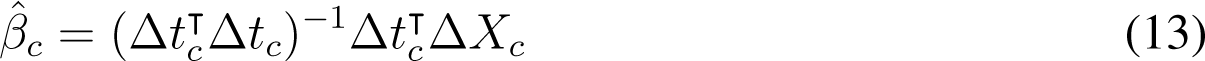

RNA velocities are calculated separately for times assigned to the response to rapamycin program and in the cell cycle program.

#### 4.6.3 Decay Rate Bounded Estimate

Estimates of decay constant are made separately for each gene *i*. Plotting *^dxi^/dt* against *x_i_* defines a convex region, and if we hold *α_i_ ≥* 0, then *−λ_i_* is the maximum slope of the line through the origin which does not intersect the convex region. *λ_i_* in this case is a minimum estimate of decay constant, constrained by the *α_i_≥* 0 requirement.

To prevent this estimate from being outlier-driven, cells are ordered by *^dxi^/dt* to *x_i_* ratio. The smallest 0.05 quantile of cells are retained and used to regress *^dxi^/dt* against *x_i_* by ordinary least-squares regression, estimating *−λ_i_* which satisfies *α_i_ ≥* 0 for the large majority of cells, while allowing some tolerance for outliers.

These decay rate estimates were then made repeatedly for cells on a sliding window, to make a time-dependent bounded estimate of mRNA decay rate.

### 4.7 Regulatory Network Modeling

Gene expression (**Z** *∈* Z*^n^*^x^*^g^|* **Z** *≥* 0) is standardized by scaling each cell *n* so that the sum of all genes *g* for the observation is equal to 3099 counts (the median count depth for all cells in the dataset). Features are then standardized by scaling each gene *g* without centering. Genes are scaled to interquartile range (IQR; Q_3_ *−* Q_2_) if the IQR^2^ is greater than gene *g* variance, and scaled to 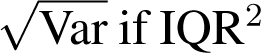 is less than gene *g* variance (Eq. 14). Any genes *g* which have a variance of zero are removed. The standardized gene expression data (**X** *∈* R*^n^*^x^*^g^|* **X** *≥* 0, Var(**x***_g_*) *≤* 1) is used for modeling.

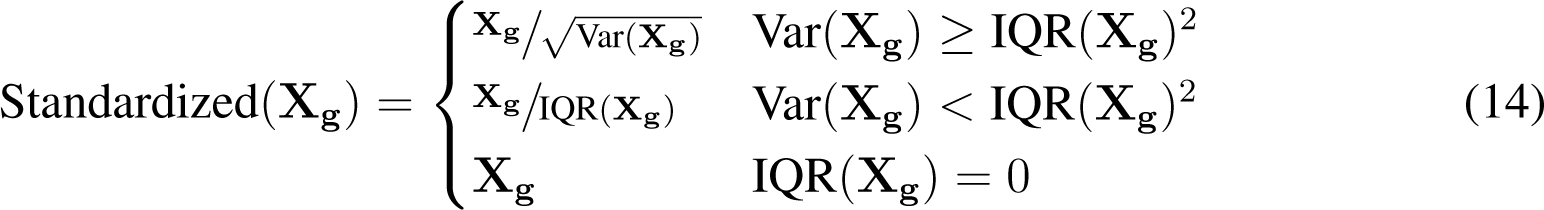

#### 4.7.1 Prior Network Knowledge

An existing gold standard regulatory network (*23*) is merged with experimental evidence from a systematic dynamic perturbation screen of yeast TFs (*62*) to generate a prior knowledge matrix of high-confidence interactions. This prior knowledge network **P** (**P** *∈ {*0, 1*}*) consists of 2799 regulatory edges, connecting 203 regulatory TFs to 1573 yeast genes.

#### 4.7.2 Linear Network Inference Model

TF Activity **A^^^** (**A^^^** *∈* R*^n^*^x^*^k^*) of TFs *k* are inferred with a linear model from expression **X** and prior knowledge **P** (Eq. 15). **A^^^** is then used to solve for gene regulatory network ***β***, regularized with least absolute shrinkage and selection operator (LASSO) (Eq. 16).

The *£*_1_ hyperparameter *λ_£_*is selected by Stability Approach to Regularization Selection (StARS) (*63, 64*). In short, the LASSO regression is sampled equally without replacement into 20 subregression problems, solving for ***β****_s_* (**s** *∈ {*1*, …,* 20*}*). For each subregression problem *s*, ***β****_s_* is calculated stepwise for decreasing values of *λ_£_*. As *λ* decreases, ***β****_s_* decreases in sparsity, going from highly sparse at large values of *λ_£_* to highly dense at small values. We select *λ* which maximizes sparsity of the subnetworks ***β****_s_* while having less than a 0.05 fraction of divergent edges (edges present in only some subnetworks and not others).

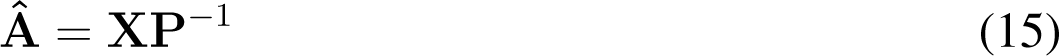

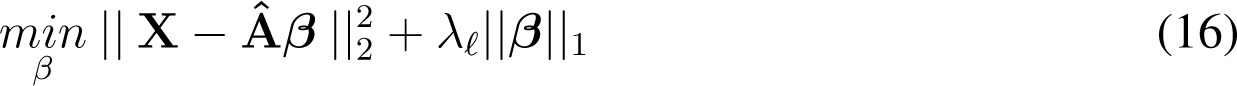

Non-zero values of ***β****_s_* are considered to be evidence for TF to gene regulation, and are ranked by explained relative variance (Eq. 24) to construct a GRN.

#### 4.7.3 Static Deep Model

For each cell (observation) *n*, expression of genes *g* (**n** *∈* R*^g^*) are linearly embedded with weights **W***_E_* into a hidden layer (**h**^(^*^T^ ^F^* ^)^ *∈* R*^k^*) with *k* nodes, representing TFs. Encoder weights **W***_E_* are pruned using prior network knowledge **P**, so that edges which have a value of 0 in **P** are set to 0 in **W***_E_* before and during training. **h**^(^*^T^ ^F^* ^)^ is constrained to positive values with a rectified linear unit (ReLU) activation function and interpreted as transcription factor activity (TFA) (Eq. 17).

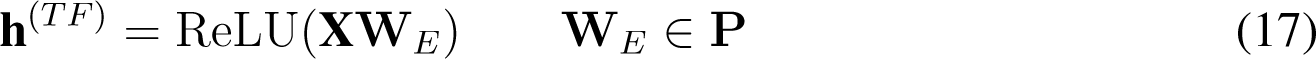

The TF hidden layer (**h**^(^*^T^ ^F^* ^)^) is linearly embedded into an intermediate hidden layer with *k* nodes (**h**(^2^) *∈* R*^k^*) with fully-connected weights **W**_2_ and a ReLU activation function (Eq. 18). **h**(^2^) is then connected to output gene expression nodes *g* with linear weights **W***_D_* (Eq. 19). If this output is count data, ReLU activation function is applied to the output nodes; if it is velocity data, no activation function is applied.

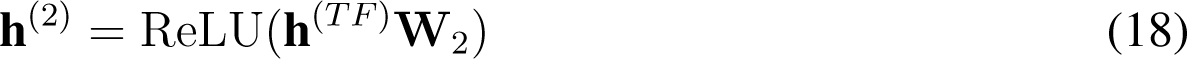

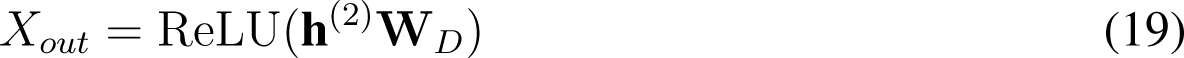

#### 4.7.4 Dynamical Deep Model

Each cell is positioned on a trajectory *τ* with length *£_t_* (*τ ∈* R*^£t^*^x^*^g^*). Trajectories *τ* are embedded into a TF hidden layer (**h**^(^*^T^ ^F^* ^)^) through a constraint mask, as for the static model. This TFA layer (**h**^(^*^T^ ^F^* ^)^) is used as input into a fully connected recursive neural network layer (**h**(^2^)) with *k* nodes at time *t*.

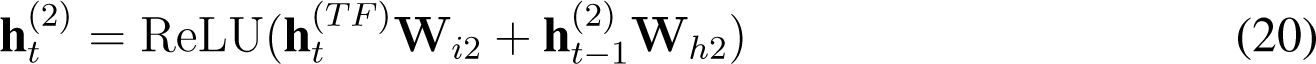

The outputs **h**(^2^) are then connected to output gene expression nodes *n* with linear weights **W***_D_*. If this output is count data, ReLU activation function is applied to the output nodes; if it is velocity data, no activation function is applied.

#### 4.7.5 Decay Model

Each cell is positioned on a trajectory *τ* with length *£_t_* (*τ ∈* R*^£t^*^x^*^g^*). Trajectories *τ* are linearly embedded with weights **W***_E_*(_1_) into a hidden layer (**h**(^1^) *∈* R^50^), a tanh activation function is applied, and **h**(^1^) is linearly embedded with weights **W***_E_*(_2_) into a second hidden layer (**h**(^2^) *∈* R^50^), followed by a Softplus activation function (Eq. 21).

**h**(^2^) is input into a fully connected recursive neural network layer (**h**(^3^)) with 50 nodes at time *t*. **h**(^3^) is connected to output decay rate nodes *g* with linear weights **W***_D_* and a Softplus activation function. Output decay rates are multiplied by -1 and then by expression of genes *g* to yield as model output RNA velocity which is a product of mRNA decay.

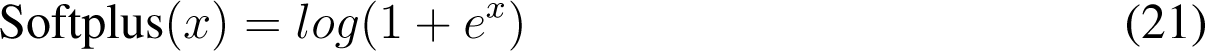

#### 4.7.6 Biophysical Model

Each cell is positioned on a trajectory *τ* with length *£_t_* (*τ ∈* R*^£t^*^x^*^g^*). This is used as input into decay model module, which yields RNA velocity as a product of mRNA decay. It is also used as input into a dynamical deep model which yields RNA velocity as a product of mRNA transcription, and is interpretable as a transcriptional gene regulatory network.

The transcriptional model is constrained to positive RNA velocity with a Softplus activation function, and the decay model is constrained by design to negative RNA velocity. The output from these two models is added to yield the biophysical RNA velocity model output.

#### 4.7.7 *In silico* TF perturbation

After calculating the TF hidden layer **h**^(^*^T^ ^F^* ^)^, **h**^(^*^T^ ^F^* ^)^ is multiplied by a diagonal perturbation matrix (**D** *∈* R*^k^*^x^*^k^*) and then used as input into the next model layer. For unperturbed predictions, **D** = **I**. For perturbed predictions, diagonal elements of **D** are 1 if the TF is included and 0 if the TF is deleted.

#### 4.7.8 Training

All models are implemented in pytorch (*65*) and trained to minimize mean squared error (Eq. 22) with batched gradient descent and the Adam optimizer. All models are trained on the CPU of a single 40-core node (dual-socket Intel Xeon Gold 6148 2.40GHz). 25% of observations from gene expression **X** are held out as validation data **X**_VALIDATION_, and the remaining 75% of observations are used as training data **X**_TRAIN_. Models are regularized by random dropout of 50% of input genes *g*, randomized for each training batch and epoch. Models trained for hyperparameter search are trained for 200 epochs. Full models used for prediction are trained for 2000 epochs.

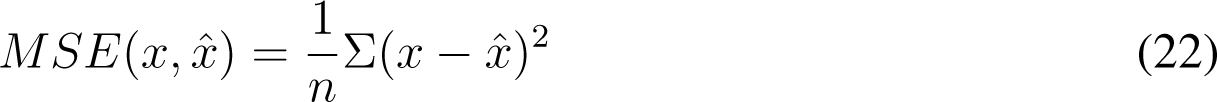

#### 4.7.9 Static Deep Model Training

Observations are sampled at random from gene expression **X**_TRAIN_ into batches of 250 observations.

#### 4.7.10 Dynamical Deep Model Training

Observations from gene expression **X**_TRAIN_ are binned by time into bins (width = 1 minute). Times are response to rapamycin **t***_RAPA_* (**t***_RAPA_ ∈* R*^n^| −* 10 *≤* **t***_RAPA_ ≤* 60) or cell cycle **t***_CC_* (**t***_CC_ ∈* R*^n^|* 0 *≤* **t***_CC_ ≤* 88). Trajectories *τ* of length *£* are constructed by randomly selecting observations from *£* sequential bins. Trajectories *τ* are then sampled at random into batches of 20 observations. After every training epoch, trajectories are discarded and new trajectories are built by randomly sampling observations into trajectories again.

#### 4.7.11 Decay Model Training

An estimate for transcript decay rate is made for every cell (See Section: Decay Rate Bounded Estimate) and multiplied by denoised count data to get **<u>^X^ X</u>** *t decay ∈* R*^n^*^x^*^g^*), which is an estimate for the negative, decay component of velocity. 25% of these observations are held out for validation, and the other 75% are used to train the decay model.

#### 4.7.12 Biophysical Model Training

The transcriptional module of the biophysical model is initialized randomly, and the decay model is initialized with weights from a pre-trained decay model. The decay model is then frozen for 100 epochs, and the transcriptional model is trained, minimizing total model loss to RNA velocity. After 100 epochs, the decay model is unfrozen and the model is trained to minimize total model loss to RNA velocity and to minimize decay model loss to the estimated negative decay component of velocity.

#### 4.7.13 Hyperparameter Tuning

Model parameters are tuned on both causal network performance and on coefficient of determination. 25% of the observations are held out of model training as validation to calculate R^2^ on data not included in training. 25% of the prior network knowledge genes are held out of the prior knowledge **P** as validation to calculate AUPR on network knowledge not included in training.

Input dropout rate is fixed to 50%, model structure is fixed, and batch sizes are fixed. Gradient descent parameters learning rate (*γ ∈ {*10*^−^*^2^, 5 *·* 10*^−^*^2^, 10*^−^*^3^, 5 *·* 10*^−^*^3^, 10*^−^*^4^, 5 *·* 10*^−^*^5^, 10*^−^*^5^, 5 *·* 10*^−^*^5^, 10*^−^*^6^, 5*·*10*^−^*^6^*}*) and weight decay (*λ ∈ {*10*^−^*^2^, 5*·*10*^−^*^2^, 10*^−^*^3^, 5*·*10*^−^*^3^, 10*^−^*^4^, 5*·*10*^−^*^5^, 10*^−^*^5^, 5*·*10*^−^*^5^, 10*^−^*^6^, 5 *·* 10*^−^*^6^, 10*^−^*^7^*}*) are tuned to optimize validation AUPR and R^2^.

#### 4.7.14 Performance

Trained models are interpreted to provide evidence for biological TF to gene regulatory relationships. A TF is removed from the model to create a *tf* -reduced model. Explained relative variance (ERV) is one minus the ratio of the residual sum of squares (RSS) (Eq. 23) of the *tf* -reduced model and the full model (Eq. 24). This is repeated for each TF to calculate a genes by TFs ERV matrix (**ERV** *∈* R*^g^*^x^*^k^|* **ERV** *≤* 1).

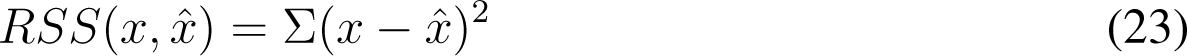

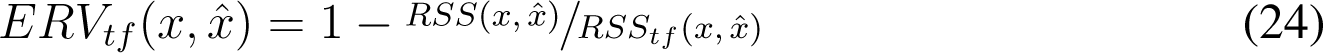

TF to gene regulatory edges are ranked by explained relative variance and compared to known TF to gene regulatory relationships derived from literature. Causal network inference performance is evaluated as Area Under the Precision - Recall curve (AUPR). Models are further evaluated based on coefficient of determination (R^2^)

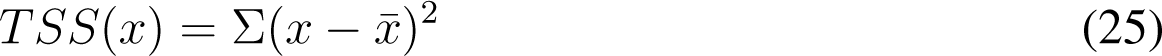

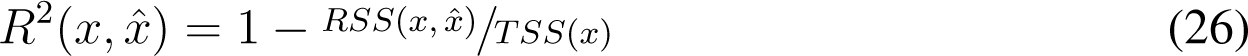

TF activity estimate performance is quantified by marginal error at the **h**^(^*^T^ ^F^* ^)^ layer, decomposing each TF’s contribution to the model loss into *L_T F_* . This is defined as the partial derivative of the model loss *L* (Mean Squared Error; Equation 22) with respect to **h**^(^*^T^ ^F^* ^)^ multiplied by **h**^(^*^T^ ^F^* ^)^ (Eq. 27), calculated using the chain rule, implemented as standard neural network backpropagation. For each trajectory, marginal error is summed over time, and the mean value of all trajectory error is reported.

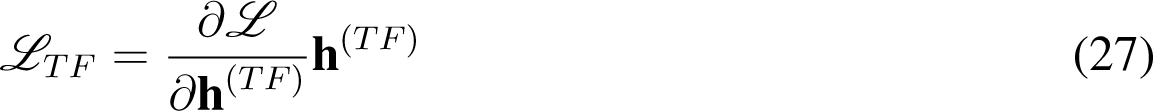

### 4.8 Visualization

All figures are generated with python using matplotlib (*46*) and inserted into this work unaltered. Schematics are drawn in Adobe Illustrator, rasterized to PNG, and imported into python with matplotlib. Data analysis and visualizations depend on the core packages numpy (*50*), pandas (*51*), scipy (*66*), scikit-learn (*67*), and scanpy (*68*).

### 4.9 Supplemental Data

- **Supplemental Data 1** is single-cell response to rapamycin count data first sequenced in this work. It is a 173348 rows × 5847 columns TSV.GZ file where the first row is a header, the first 5843 columns are integer gene counts, and the final 4 columns (’Gene’, ‘Replicate’, ‘Pool’, and ‘Experiment’) are cell-specific metadata.
- **Supplemental Data 2** is bulk response to rapamycin count data first sequenced in this work. It is a 33 rows × 5847 columns TSV.GZ file where the first row is a header, the first 5843 columns are integer gene counts, and the final 4 columns (’Oligo’, ‘Time’, ‘Replicate’, and ‘Sample barcode’) are sample-specific metadata.
- **Supplemental Data 3** is single-cell count data published as GSE125162 and re-analyzed with the pipeline used for single-cell quantification in this work. It is a 65068 rows × 5850 columns TSV.GZ file where the first row is a header, the first 5843 columns are integer gene counts, and the final 7 columns (’Condition’, ‘Sample’, ‘Genotype Group’, ‘Genotype Individual’, ‘Genotype’, ‘Replicate’, ‘Cell Barcode’) are cell-specific metadata.
- **Supplemental Data 4** is the four deep learning models trained in this work. It is a TAR.GZ file containing the final biophysical transcription/decay model, the pre-trained decay model, the velocity prediction model, and the count prediction model. Each model file is an h5 file containing a pytorch model that can be loaded with supirfactor dynamical.read().
- **Supplemental Data 5** is the prior knowledge network used to constrain the models for TF interpretability. It is a 1574 rows × 204 columns [Genes x TFs] TSV.GZ file where the first row is a header with TF names, the first column is an index of gene names, and TF-gene interactions are indicated by non-zero values in the matrix. There are 2799 TF-gene interactions.
- **Supplemental Table 6** is the oligonucleotide sequences used in this work. It is a TSV file with a header row.
- **Supplemental Table 7** is the yeast strains used in this work. It is a TSV file with a header row.
- **Supplemental Table 8** is gene metadata used in this work (e.g. Ribosomal Protein gene labels, etc). It is a TSV file with a header row.
- **Supplemental Table 9** is FY4/5 growth curve data generated in this work. It is a 20 rows × 7 columns TSV file where the first row is a header with replicate IDs, the first column is an index of times in minutes, and values are cell densities in YPD culture, in units of 10^6^ cells / mL.
- **Supplemental Data 10** is a TAR.GZ file containing the yeast SacCer3 genome, modified to add UTR sequences, that was used to generate transcripts for kallisto pseudoalignment in this work.

## Acknowledgements

We thank past and present members of the Gresham and Bonneau labs for discussions and valuable feedback on this manuscript. We also thank the staff of the Flatiron Institute Scientific Computing Core for their tireless efforts to build and maintain the High Performance Computing resources which we rely on. This work was supported in part through the NYU IT High Performance Computing resources, services, and staff expertise.

## 4.11 Author Contributions

Conceptualization: CAJ, AT, RB, DG; Methodology: CAJ, MBA, AT; Software: CAJ, MBA, AT; Validation: CAJ, MBA, IS, ASH; Formal analysis: CAJ, MBA; Investigation: CAJ, MBA, IS, ASH; Resources: CAJ, IS, ASH; Data curation: CAJ, MBA; Visualization: CAJ, MBA; Writing – original draft preparation CAJ, MBA, AT, RB, DG; Writing – review and editing: CAJ, MBA, AT, RB, DG; Supervision: CAJ, RB, DG; Funding acquisition: RB, DG; Project administration: CAJ, RB, DG

## 4.13 Data and materials availability

Sequencing data has been deposited in NCBI GEO under accession GSE242556. Code to assign cells to times and to calculate RNA velocity is available on GitHub and PyPi as the inferelator-velocity package. Code for dynamical deep learning modeling is available on GitHub and PyPi as the supirfactor-dynamical package. Code to generate all figures used in this manuscript is available on GitHub. Strains generated for this work are available upon request.

---

**Supplemental Figure 1:**
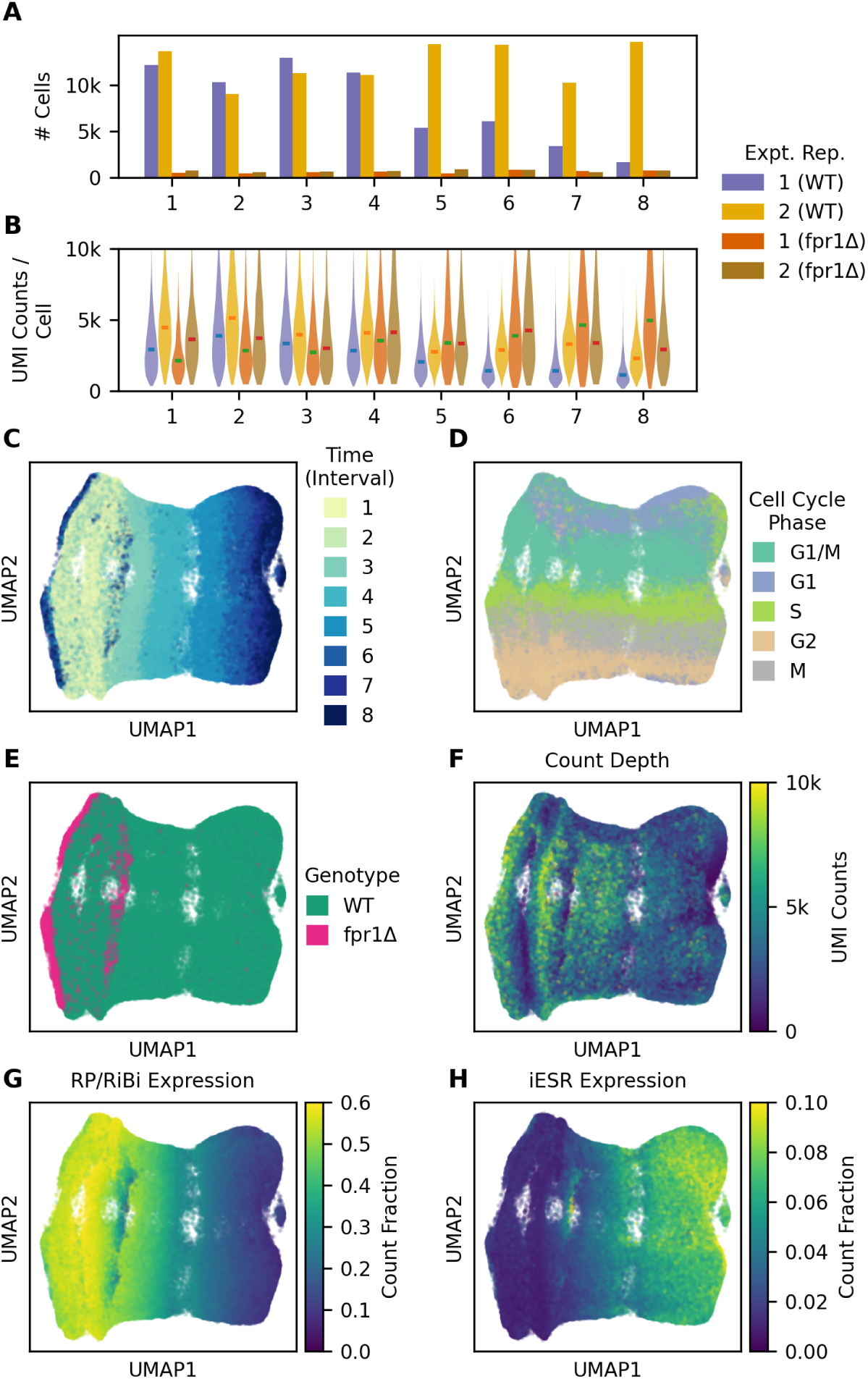
Summary of *Saccharomyces cerevisiae* single-cell transcriptomics (**A**) Number of wild-type (WT) and rapamycin non-responsive (fpr1Δ) cells captured in each sampling interval for each experimental replicate. (**B**) Distribution of the number of counts after processing Unique Molecular Identifiers (UMIs) in each sampling interval, experimental replicate, and yeast genotype. (**C**) 2D UMAP plot of single-cell *Saccharomyces cerevisiae* responding to rapamycin treatment, colored by sampling time, reproduced from Figure 1C. (**D-H**) 2D UMAP plot colored by estimated cell-cycle phase (**D**), yeast genotype (**E**), number of counts per cell (**F**), Ribosomal protein (RP) and Ribosomal biogenesis (RiBi) gene expression as a proportion of total gene expression (**G**), and Induced Environmental Stress Response (iESR) gene expression as a proportion of total gene e_5_x_2_pression (**H**).

**Supplemental Figure 2:**
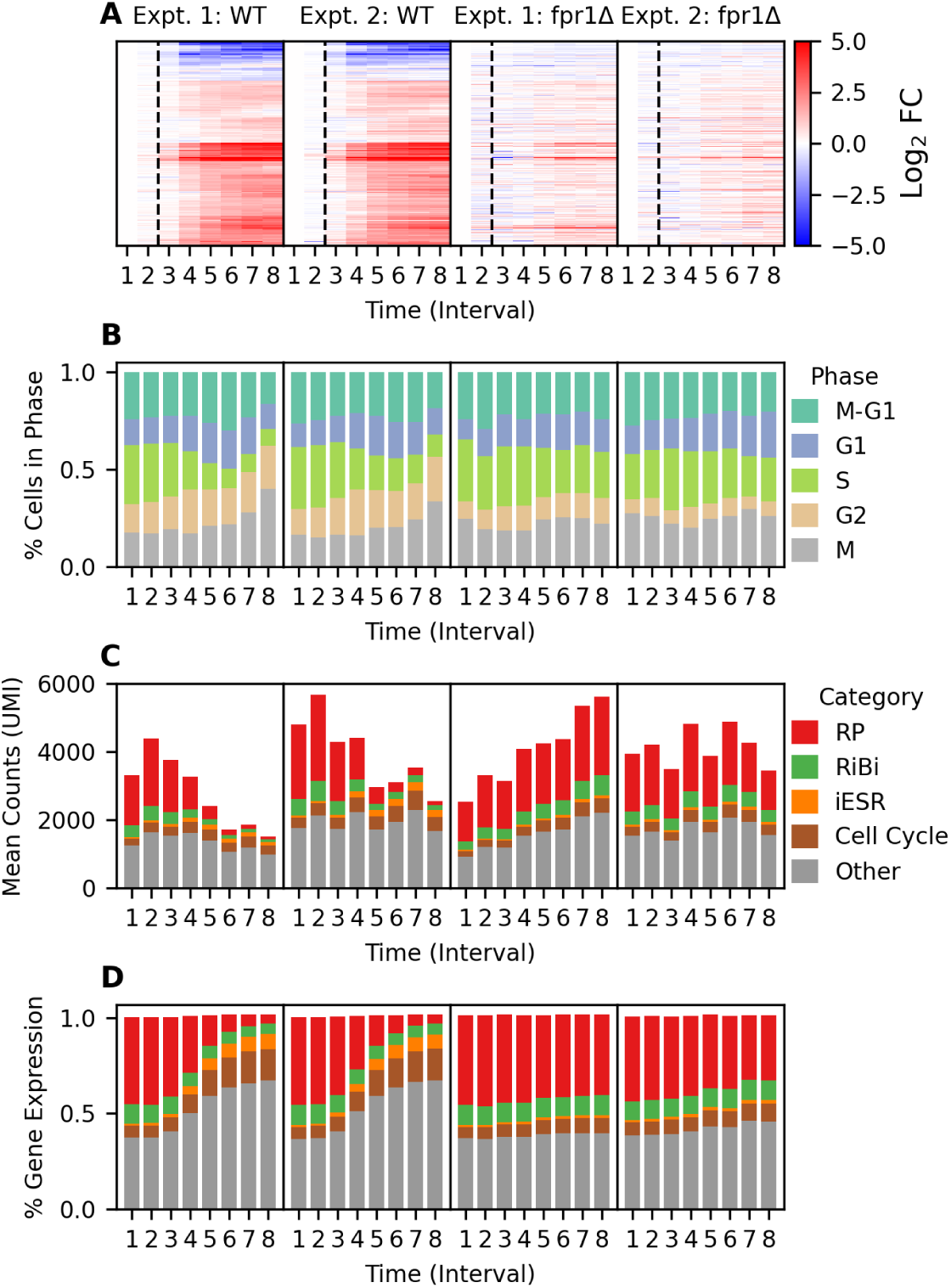
Summary of *Saccharomyces cerevisiae* single-cell changes between sampling intervals and experimental replicates (A) Differential expression (Log_2_ Fold Change compared to untreated sampling interval 1 of yeast transcriptome in wild-type (WT) and rapamycin non-responsive (fpr1Δ) cells in each experimental replicate. (B) Proportion of the number of cells in each estimated cell-cycle phase between sampling intervals and experimental replicates. (C) Median proportion of the number of counts from each category of genes between sampling intervals and experimental replicates. (D) Median absolute number of counts from each category of genes between sampling intervals and experimental replicates.

**Supplemental Figure 3:**
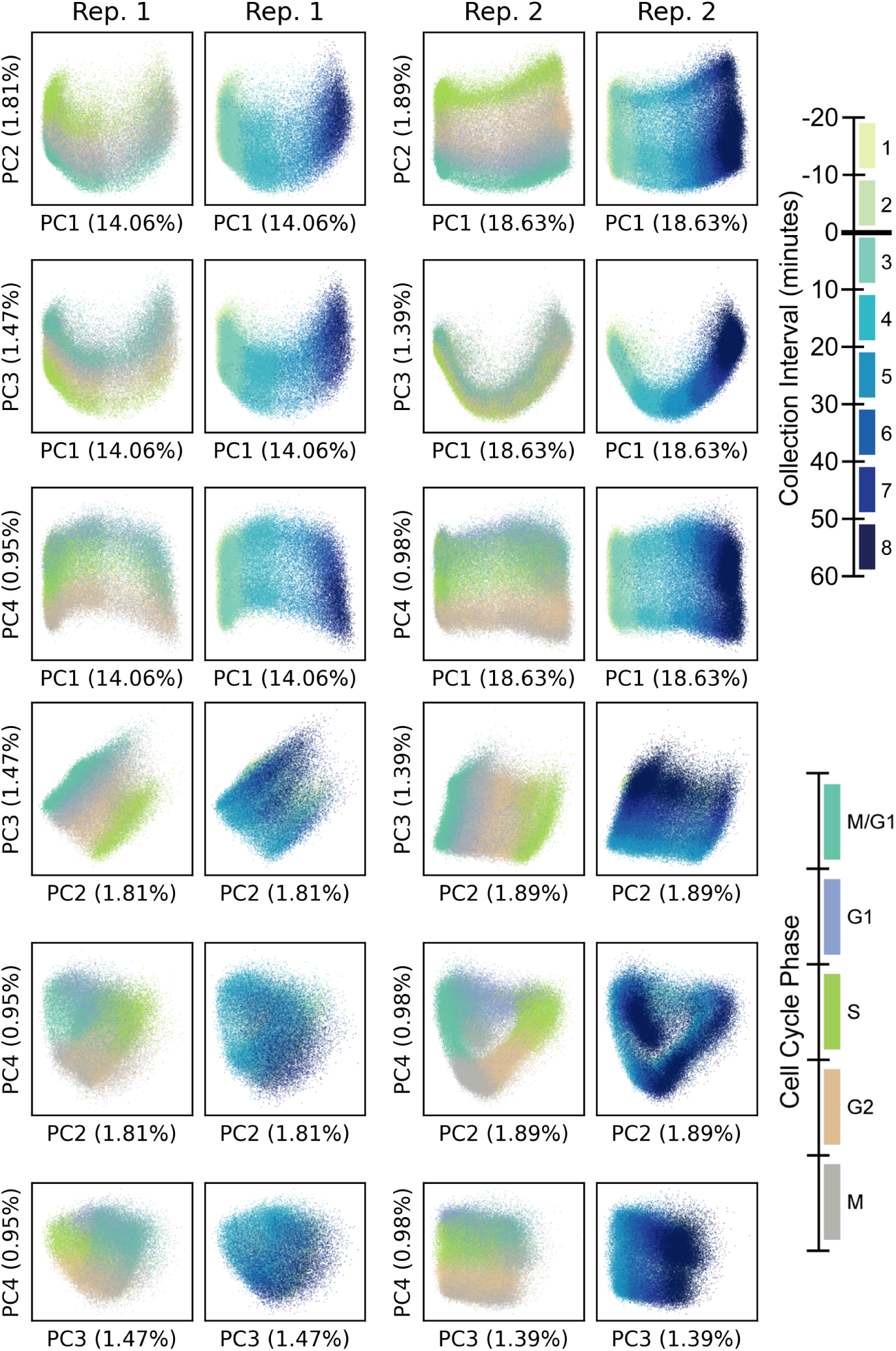
Principal component (PC) plots of the first four principal components (PC1-PC4), annotated with percent variance explained by that PC. Cells from different experimental replicates are plotted separately. Only wild-type cells are plotted. Each plot is duplicated and colored by cell cycle phase (**left**) or sampling interval (**right**).

**Supplemental Figure 4:**
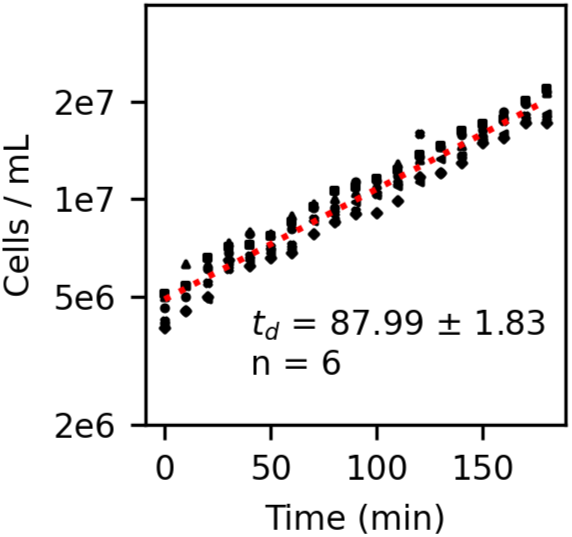
Growth rate of exponentially growing FY4/5 cells in rich YPD media. Individual cell densities for 6 replicate cultures plotted against time. Dotted line is the ordinary least-squares regression line of the form *log*_2_(*Density*) = *at* + *b*. Doubling time *t_d_* (min) is ^1^*/a* of the regression slope *±* standard error.

**Supplemental Figure 5:**
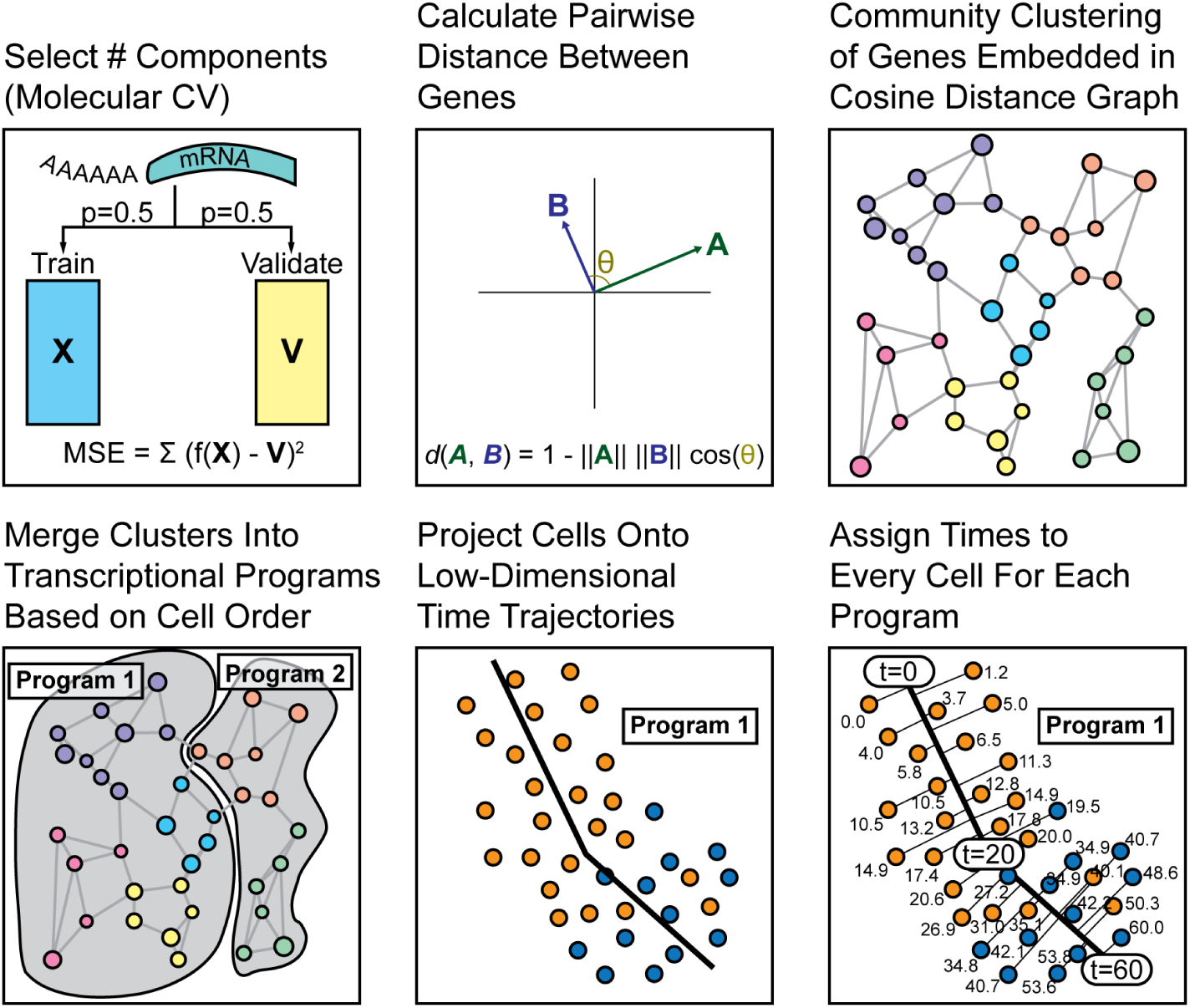
Schematic diagram of computational method for assigning cells specific times in minutes for response to rapamycin treatment and the cell cycle. **1**: Expression data is denoised by retaining the first *n* PCs, chosen by molecular cross validation, and discarding variance from other PCs. **2**: Genes are embedded into a graph using Cosine Distance as the distance metric. **3**: Genes are grouped into clusters based on the Leiden community clustering algorithm. **4**: Clusters of genes are further merged into programs based on circular rank correlation between the first principal component, calculated separately for each gene cluster. **5**: Low-dimensional time trajectories are constructed by splines between the centroids of metadata-assigned cell groups in *m* dimensional principal component space, where *m* is chosen for each program’s genes separately by molecular cross validation. **6**: Times in minutes are assigned by projecting cells onto the low-dimensional time trajectories.

**Supplemental Figure 6:**
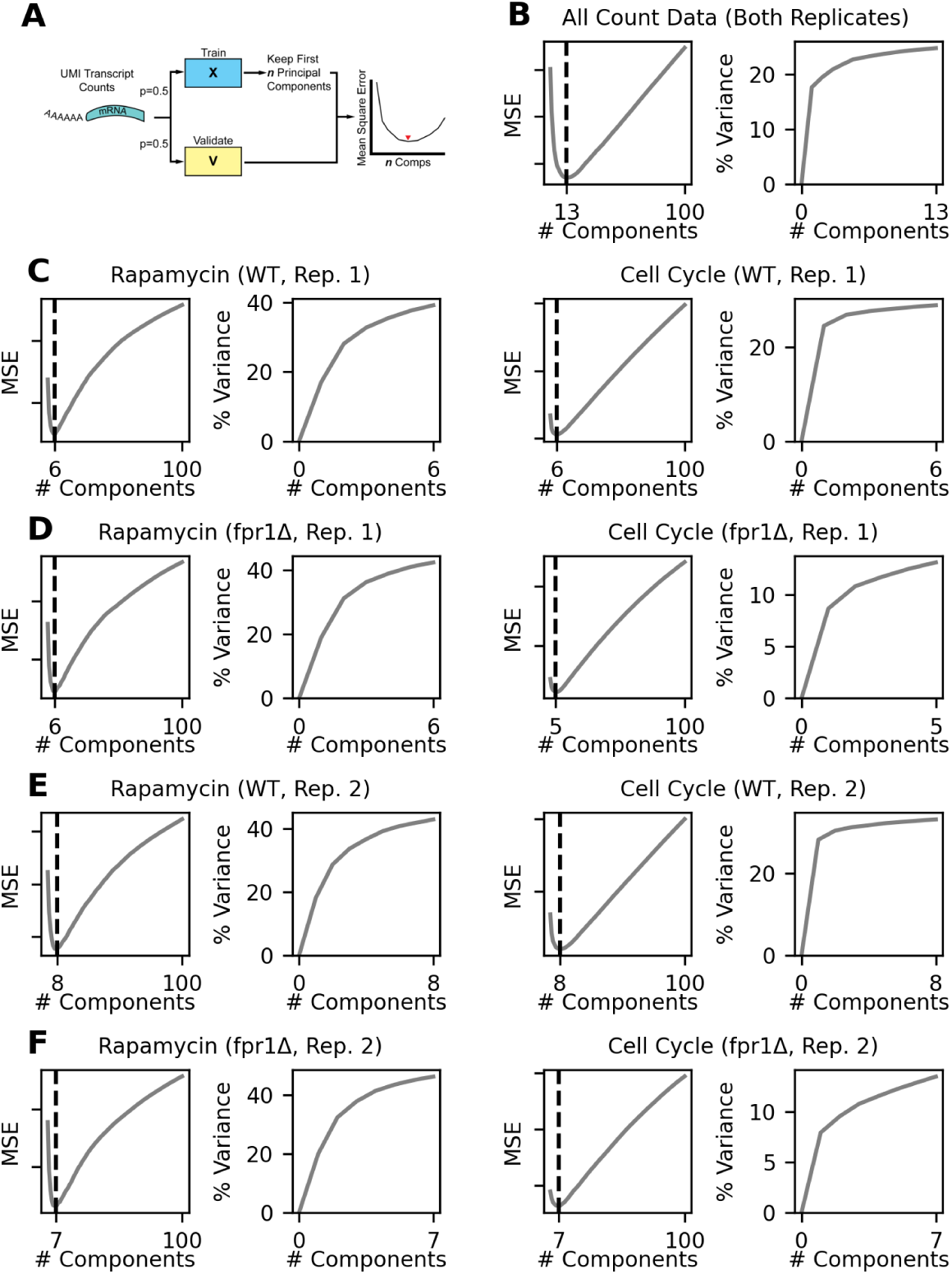
Molecular cross-validation (MCV) to select the number of principal components for gene to program assignment (**A**) MCV schematic. UMI counts are randomly partitioned into two cells by genes matrices, one of which is used for testing (X) and one of which is used for validation. Test matrix is denoised by projecting into principal component space, retaining only the top *n* principal components, and then transforming back into expression space. The optimal number of principal components minimizes mean squared error between the denoised test matrix X and the unmodified validation matrix V. (**B**) MCV results for all cells and all genes, showing mean squared error (MSE) that identifies 13 PCs as optimal, and the cumulative variance explained by 13 PCs. (**C-F**) MCV results for each program (rapaycin treatment and cell cycle) shown separately for replicate 1 wild-type (**C**), replicate 1 fpr1Δ (**D**), replicate 2 wild-type (**E**), and replicate 2 fpr1Δ (**F**).

**Supplemental Figure 7:**
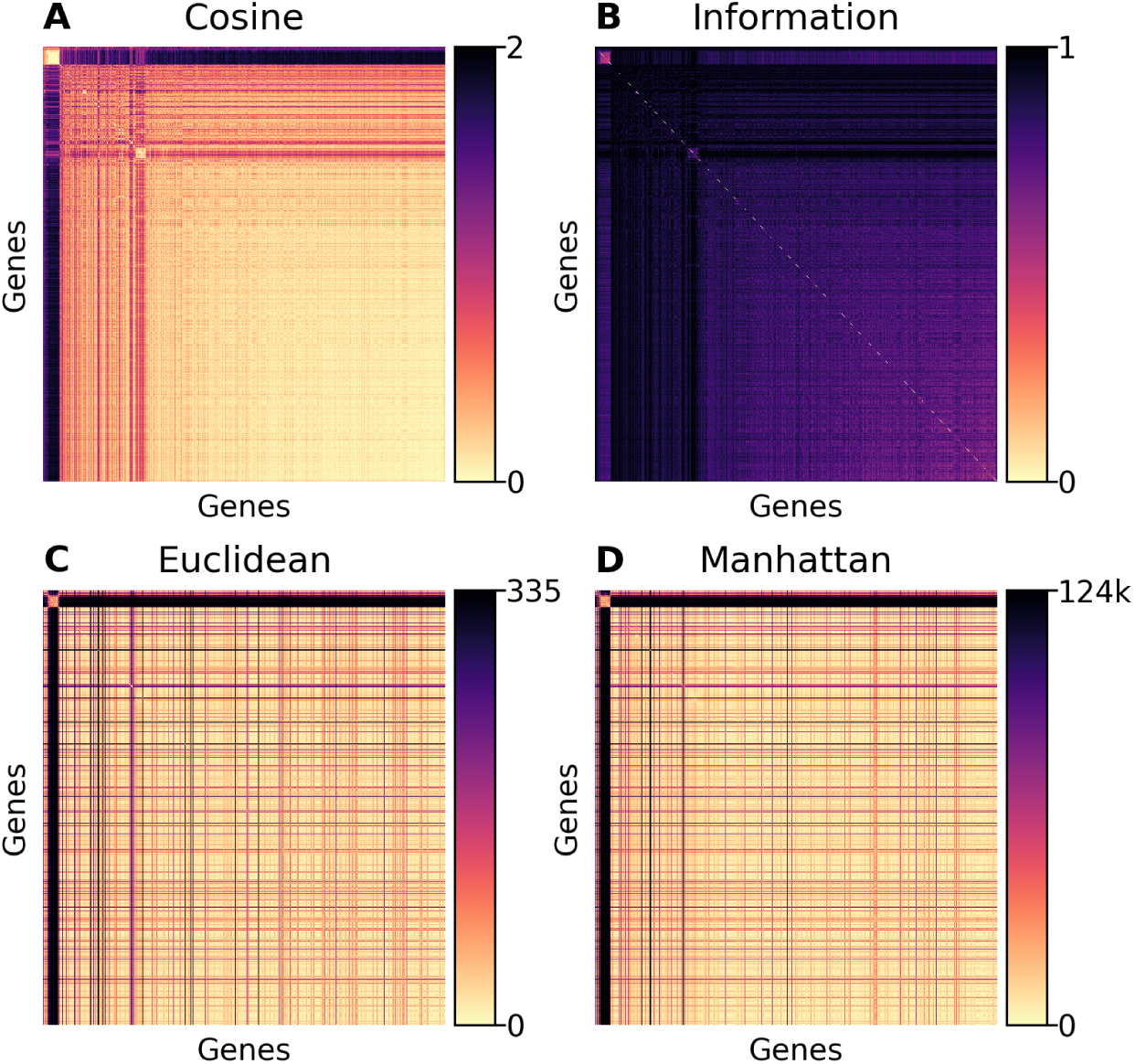
Distance metric heatmaps between genes for all cells for (**A**) Cosine distance, (**B**) Information distance, (**C**) Euclidean distance, and (**D**) Manhattan distance. Genes are ordered identically for each panel, based on hierarchical clustering using cosine distance.

**Supplemental Figure 8:**
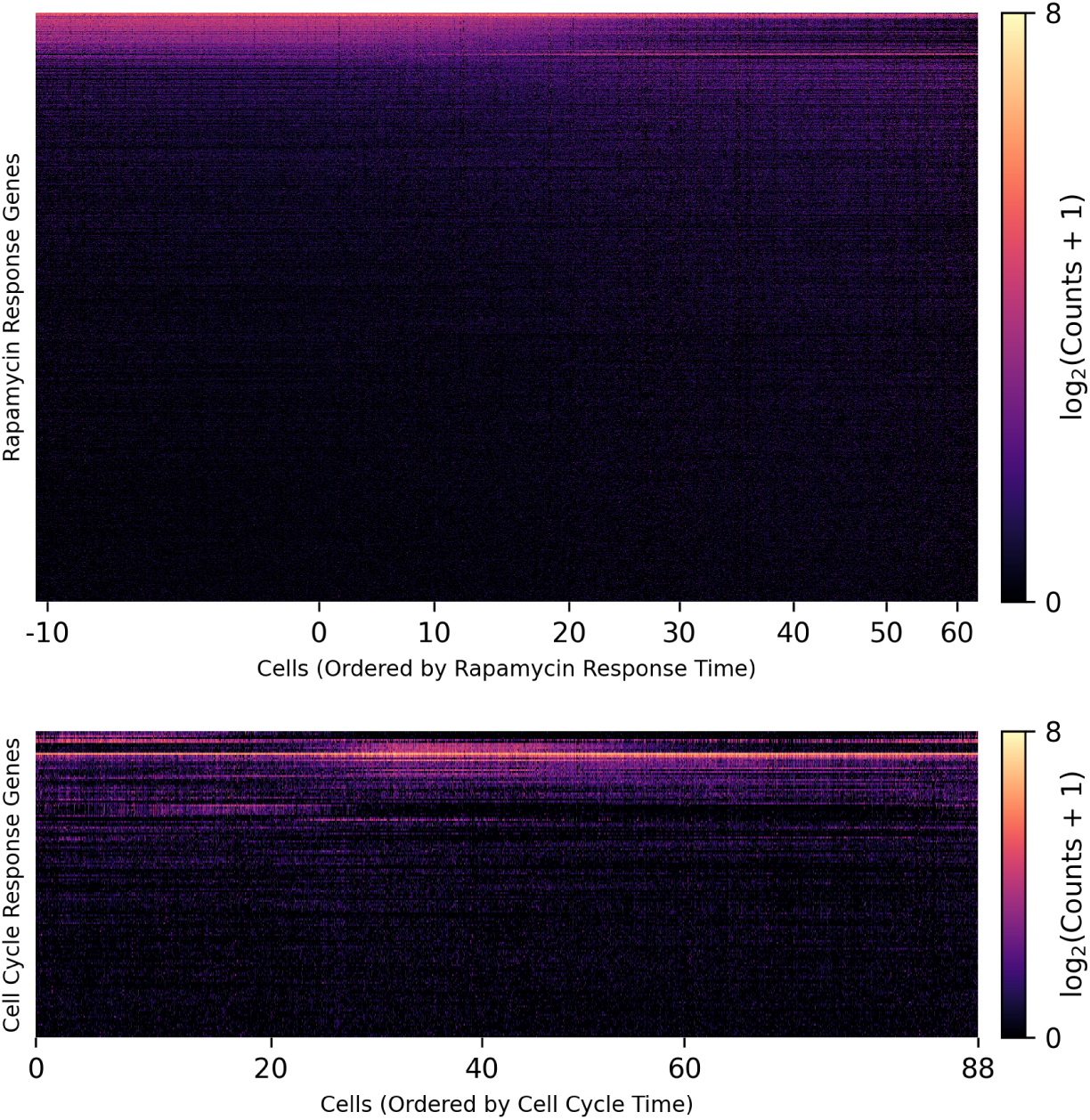
Expression heatmaps of genes for each program on the Y-axis with wild-type cells (n=162,529) on the X-axis (A) Expression of rapamycin treatment response genes (n=5379), with cells on the X-axis ordered by rapamycin response time (B) Expression of cell cycle response genes (n=348), with cells on the X-axis ordered by cell cycle time

**Supplemental Figure 9:**
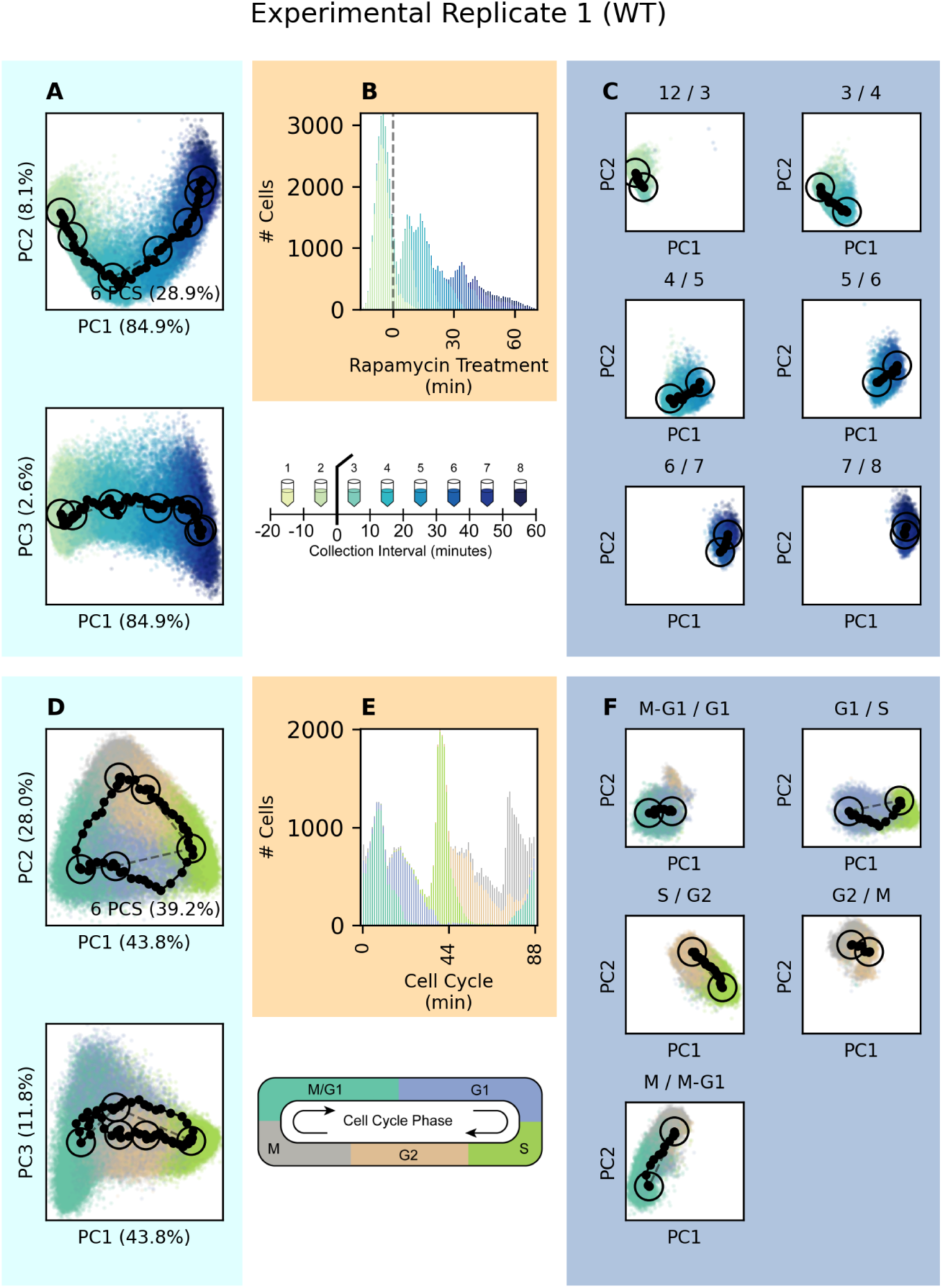
Assigning cell times to experimental replicate 1 wild-type cells. (**A**) Principal component plots of cells using only rapamycin treatment program genes. Collection time centroids are circled and connected by dashed spline. 6 PCs are used in total for time projection. (**B**) Histogram of assigned cell cycle times, colored by collection interval. (**C**) Principal component plots of cells in adjacent collection intervals, showing their centroids circled, the spline connection as a dashed line, and the shortest-walk path between them. (**D**) Principal component plots of cells using only cell cycle program genes. Cell cycle phase centroids are circled and connected by dashed spline. 6 PCs are used in total for time projection. (**E**) Histogram of assigned cell cycle times, colored by cell cycle phase. (**F**) Principal component plots of cells in adjacent phases, showing their centroids circled, the spline connection as a dashed line, and the shortest-walk path between them.

**Supplemental Figure 10:**
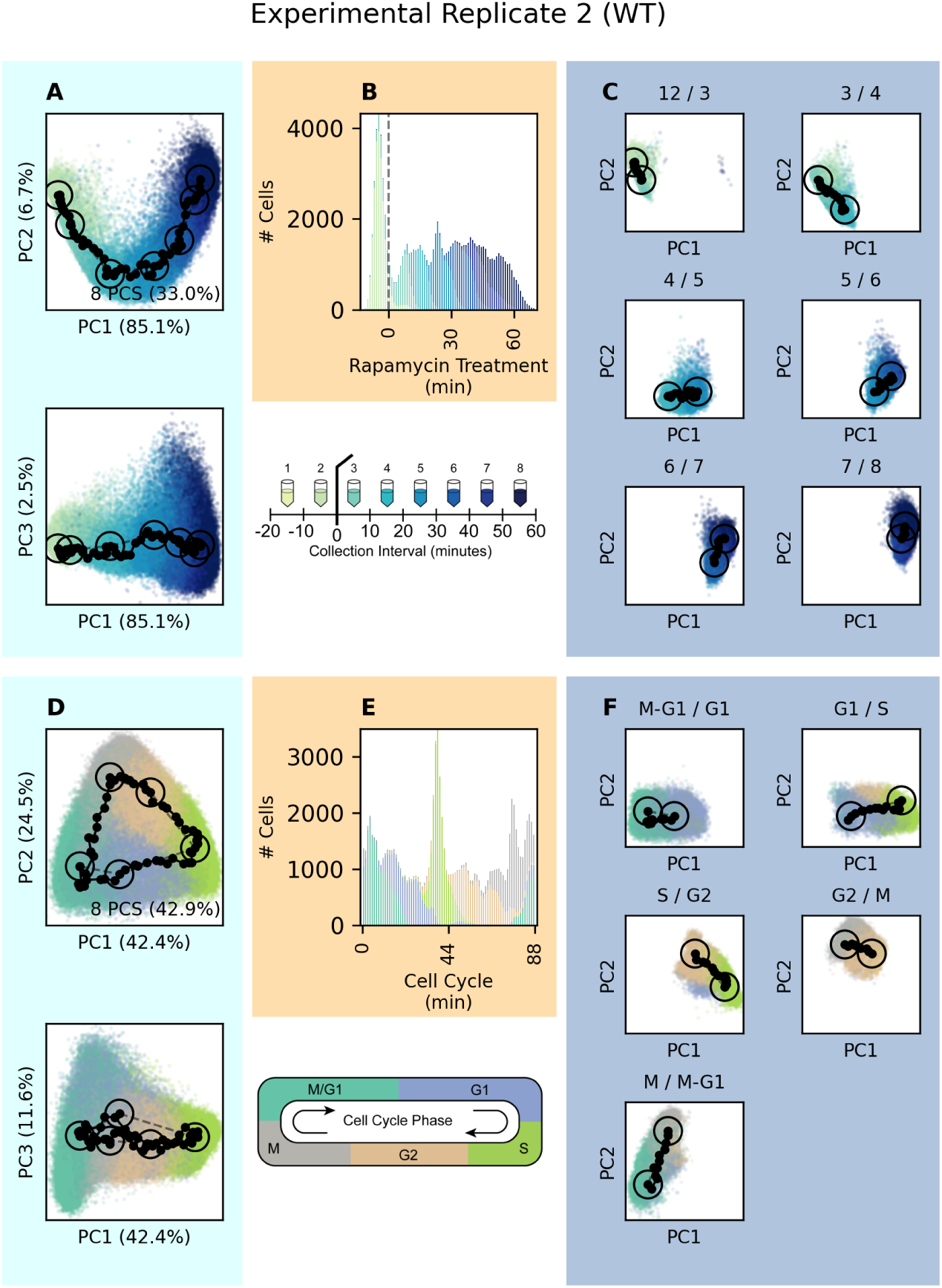
Assigning cell times to experimental replicate 2 wild-type cells. (**A**) Principal component plots of cells using only rapamycin treatment program genes. Collection time centroids are circled and connected by dashed spline. 8 PCs are used in total for time projection. (**B**) Histogram of assigned cell cycle times, colored by collection interval. (**C**) Principal component plots of cells in adjacent collection intervals, showing their centroids circled, the spline connection as a dashed line, and the shortest-walk path between them. (**D**) Principal component plots of cells using only cell cycle program genes. Cell cycle phase centroids are circled and connected by dashed spline. 8 PCs are used in total for time projection. (**E**) Histogram of assigned cell cycle times, colored by cell cycle phase. (**F**) Principal component plots of cells in adjacent phases, showing their centroids circled, the spline connection as a dashed line, and the shortest-walk path between them.

**Supplemental Figure 11:**
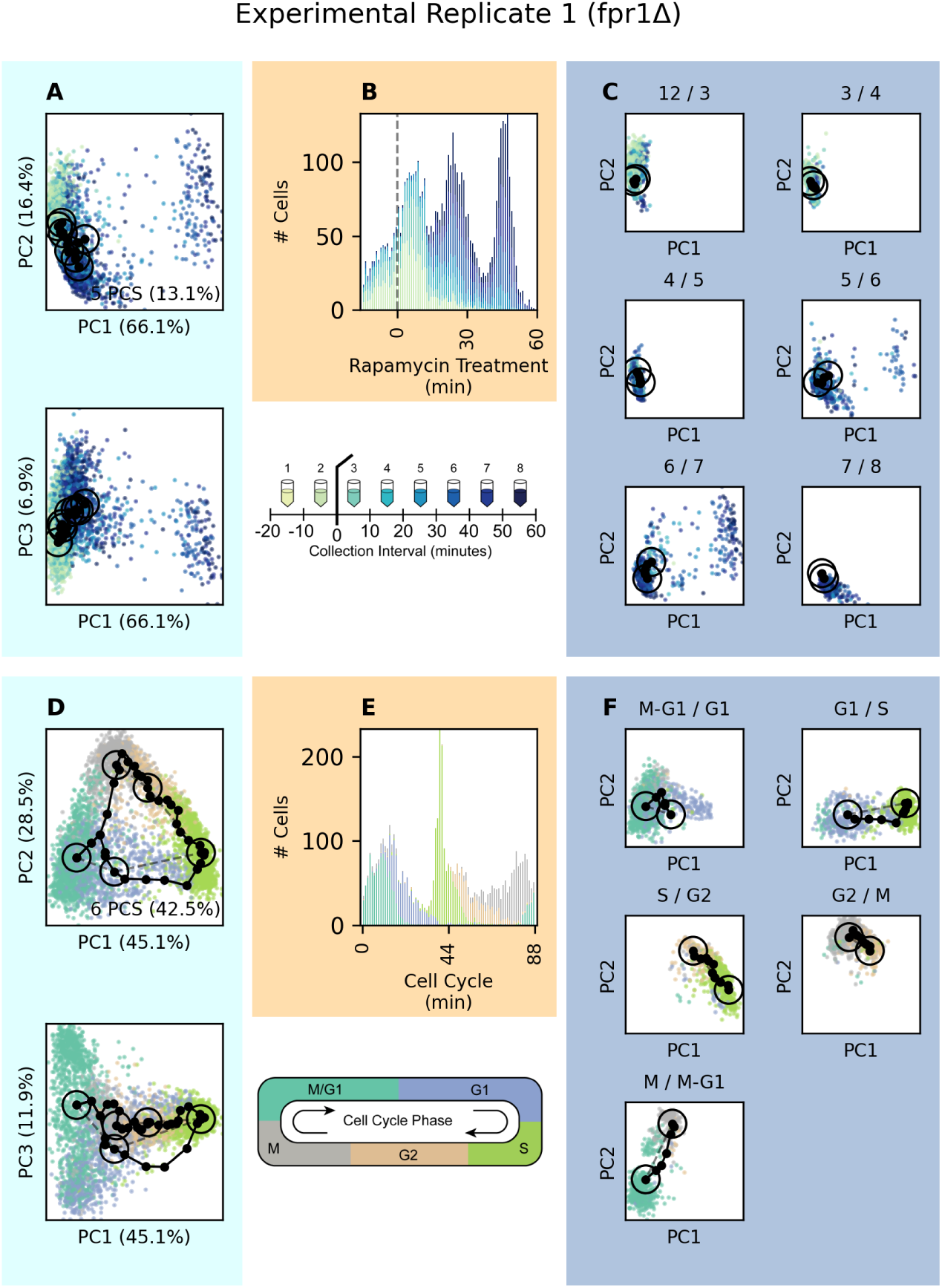
Assigning cell times to experimental replicate 1 fpr1Δ cells. (**A**) Principal component plots of cells using only rapamycin treatment program genes. Collection time centroids are circled and connected by dashed spline. 5 PCs are used in total for time projection. (**B**) Histogram of assigned cell cycle times, colored by collection interval. (**C**) Principal component plots of cells in adjacent collection intervals, showing their centroids circled, the spline connection as a dashed line, and the shortest-walk path between them. (**D**) Principal component plots of cells using only cell cycle program genes. Cell cycle phase centroids are circled and connected by dashed spline. 6 PCs are used in total for time projection. (**E**) Histogram of assigned cell cycle times, colored by cell cycle phase. (**F**) Principal component plots of cells in adjacent phases, showing their centroids circled, the spline connection as a dashed line, and the shortest-walk path between them.

**Supplemental Figure 12:**
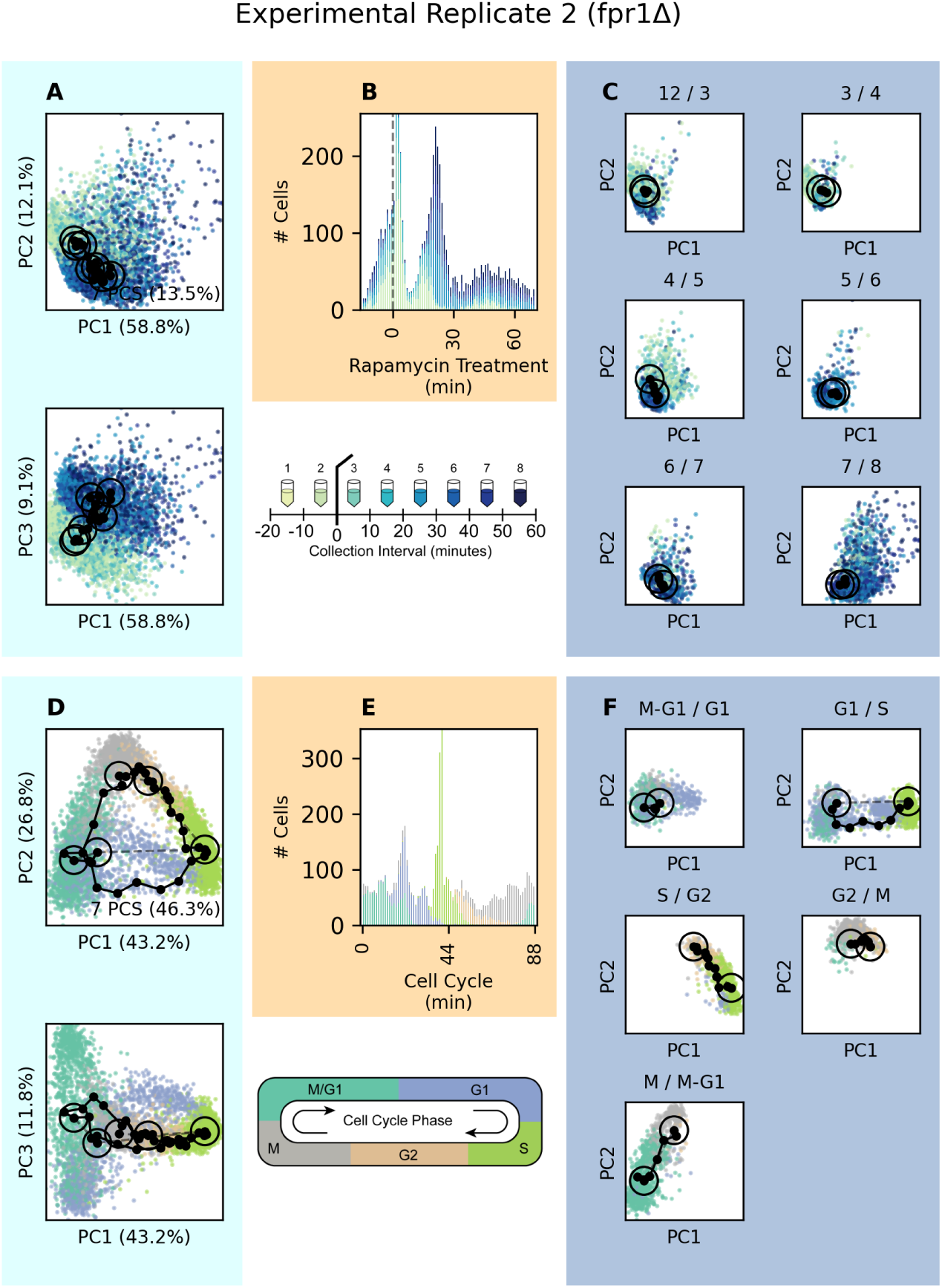
Assigning cell times to experimental replicate 2 fpr1Δ cells. (**A**) Principal component plots of cells using only rapamycin treatment program genes. Collection time centroids are circled and connected by dashed spline. 7 PCs are used in total for time projection. (**B**) Histogram of assigned cell cycle times, colored by collection interval. (**C**) Principal component plots of cells in adjacent collection intervals, showing their centroids circled, the spline connection as a dashed line, and the shortest-walk path between them. (**D**) Principal component plots of cells using only cell cycle program genes. Cell cycle phase centroids are circled and connected by dashed spline. 7 PCs are used in total for time projection. **E**) Histogram of assigned cell cycle times, colored by cell cycle phase. (**F**) Principal component plots of cells in adjacent phases, showing their centroids circled, the spline connection as a dashed line, and the shortest-walk path between them.

**Supplemental Figure 13:**
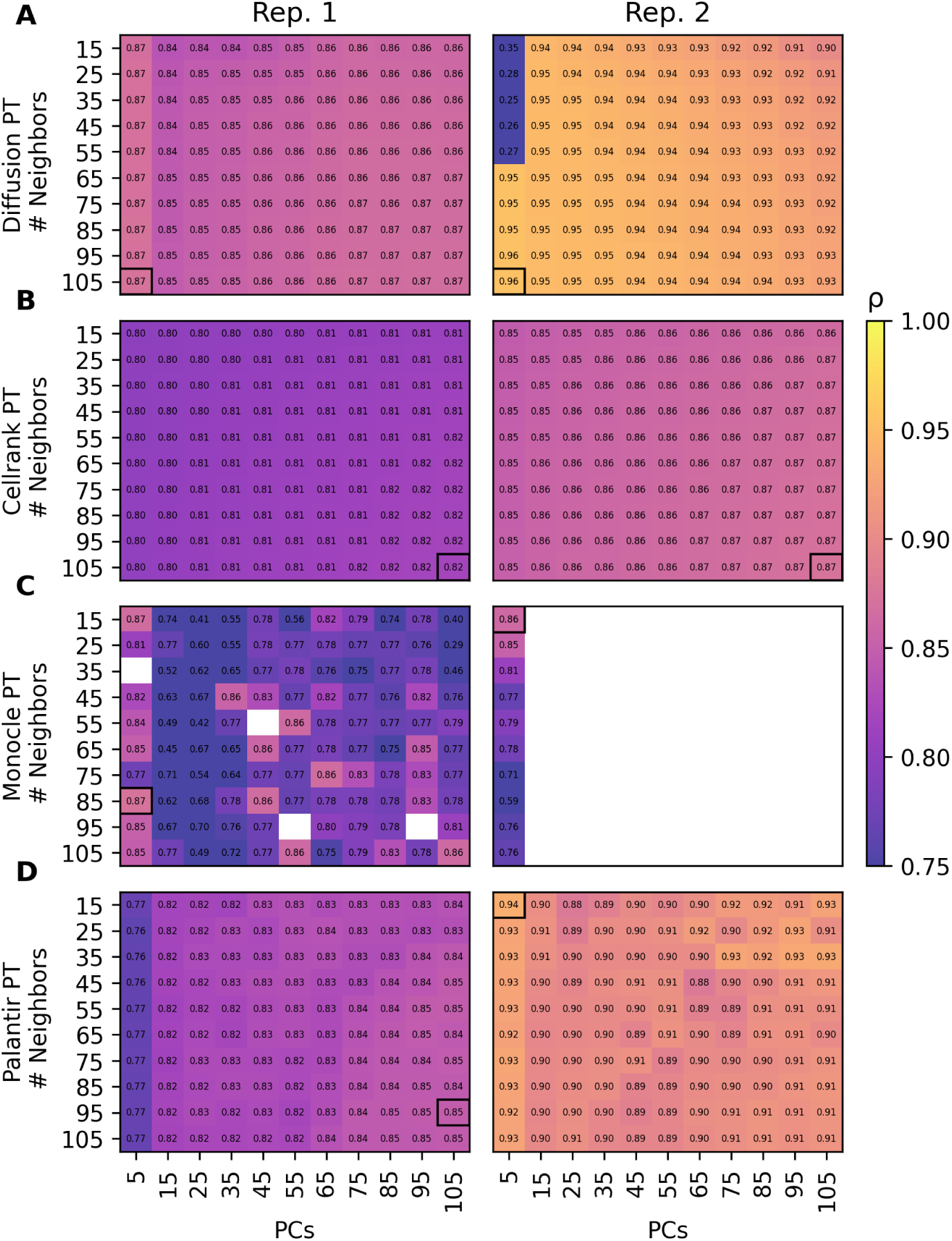
Grid search to identify optimal pseudotime *k*-nearest neighbors (Y-axis) and principal component (X-axis) hyperparameters. Scores are Spearman rho coefficients between pseudotime and the sampling interval time labels. White-background grid squares with no score indicate the presence of non-finite pseudotime values for some cells that cannot be scored. (**A-D**) Scores are reported by experimental replicate for only wild-type cells. Methods are (**A**) scanpy diffusion pseudotime where the number of diffusion components is set equal to the number of principal components, (**B**) cellrank CytoTrace pseudotime, (**C**) monocle3 pseudotime, and (**D**) palantir pseudotime where the number of diffusion components is set equal to the number of principal components.

**Supplemental Figure 14:**
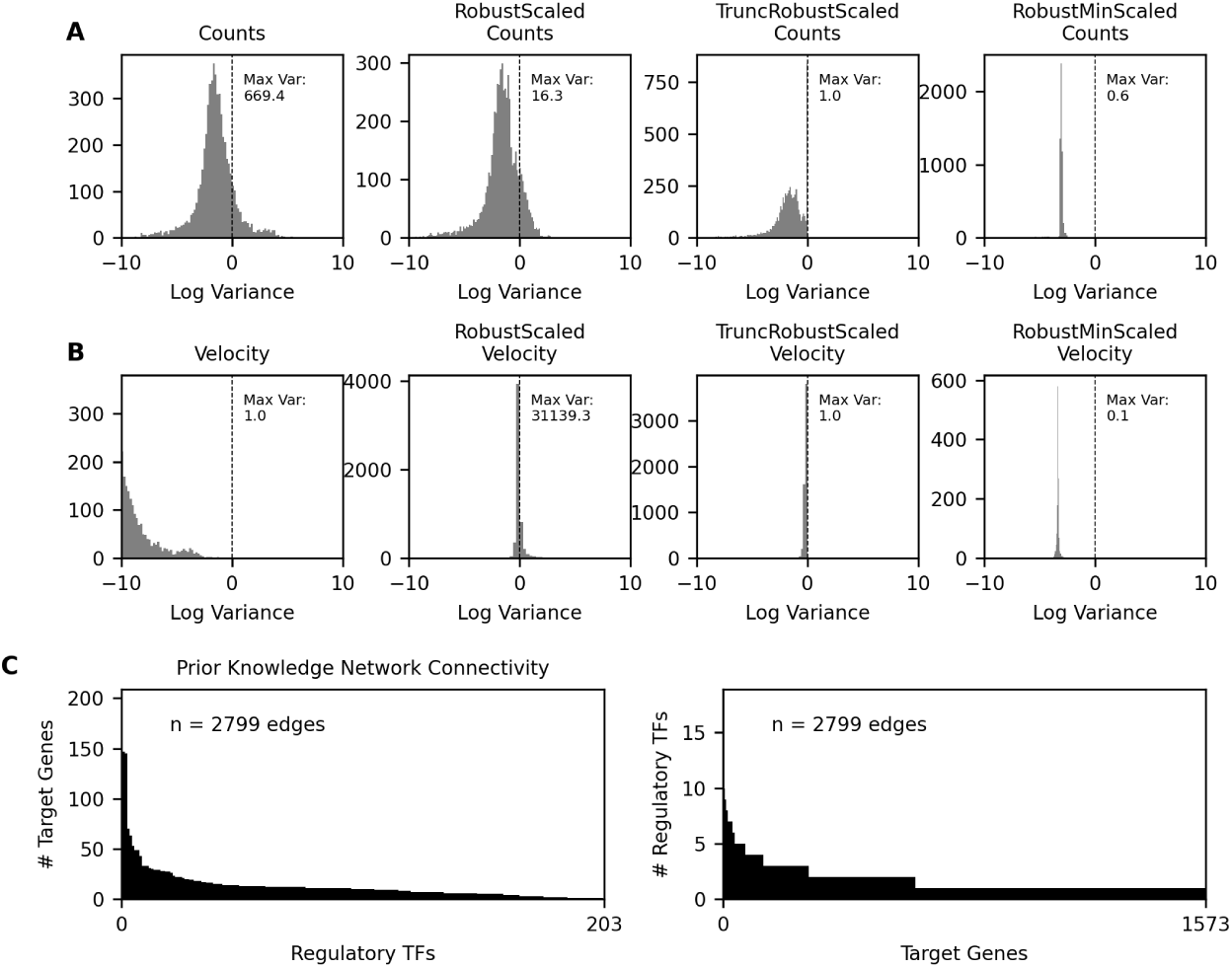
Comparison of Preprocessing and Prior Network Structure (**A**) Histograms of variance per gene in RNA expression (count) data when unscaled, when scaled to interquartile range (RobustScaled), when scaled to interquartile range with a maximum gene variance of 1 (TruncRobustScaled), and when scaled to values between 0-1 (RobustMinScaled). The maximum variance for a feature is annotated on each plot. (**B**) Histograms of variance per gene in RNA velocity (rate of change) as in **A** (**C**) Connectivity histogram of prior network knowledge between 1573 genes and 203 regulatory TFs, with 2799 regulatory network edges

**Supplemental Figure 15:**
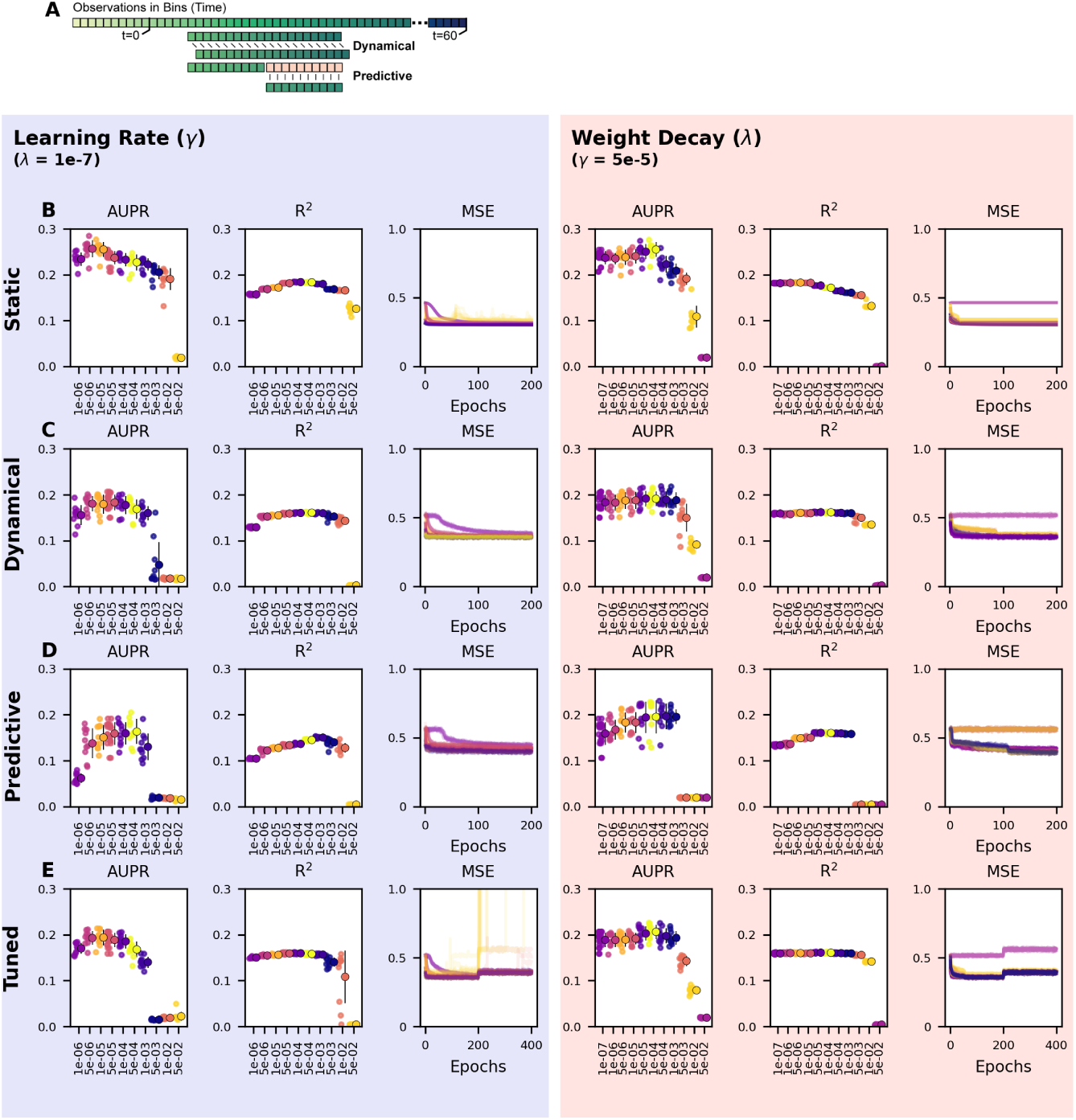
Hyperparameter search for rapamycin response regulatory network inference models. Individual cells are binned into 70 one-minute-wide bins between -10 minutes and 60 minutes, and 20 minute cell trajectories are randomly selected for each training epoch by choosing one cell from each bin. (**A**) Comparison of training strategies for Dynamical models (comparing *t* + 1 predictions only) and Predictive models (comparing a sequence of 10 predictions) (**B**) Static model performance quantified by Area Under the Precision Recall curve (AUPR), coefficient of determination (R^2^), and mean squared error (MSE) per training epoch. Performance is shown for a range of Adam optimizer learning rates (*γ*) with weight decay (*λ*) held constant, and a range of weight decays with learning rate held constant. (**C**) Dynamical model (*t* + 1 predictions) performance quantified as in **B** (**D**) Predictive model (10 sequential predictions) performance quantified as in **B** (**E**) Tuned model (training 200 epochs with the dynamical training strategy, followed by 200 epochs with the predictive model training strategy) performance quantified as in **B**

**Supplemental Figure 16:**
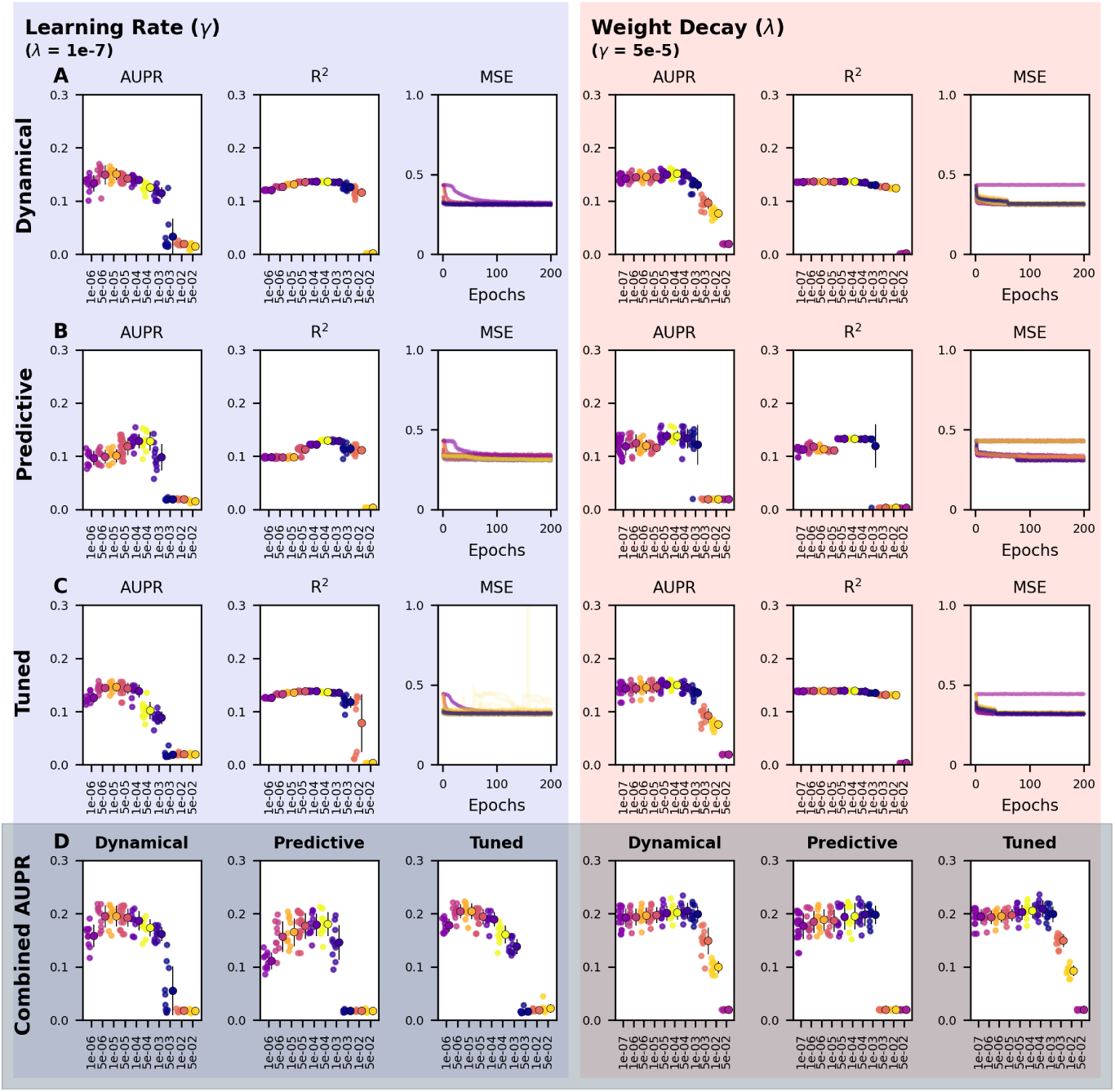
Hyperparameter search for cell cycle regulatory network inference models. Individual cells are binned into 88 one-minute-wide bins between 0 minutes and 88 minutes, and 20 minute cell trajectories are randomly selected for each training epoch by choosing one cell from each bin. (**A**) Dynamical model (*t* + 1 predictions) performance quantified by AUPR, R^2^, and mean squared error (MSE) per training epoch. Performance is shown for a range of Adam optimizer learning rates (*γ*) with weight decay (*λ*) held constant, and a range of weight decays with learning rate held constant. (**B**) Predictive model (10 sequential predictions) performance quantified as in **A** (**C**) Tuned model (training 200 epochs with the dynamical training strategy, followed by 200 epochs with the predictive model training strategy) performance quantified as in **A** (**D**) GRNs combined from rapamycin and cell cycle response models by taking the maximum explained relative variance for all regulatory edges from each model. Performance is evaluated by AUPR.

**Supplemental Figure 17:**
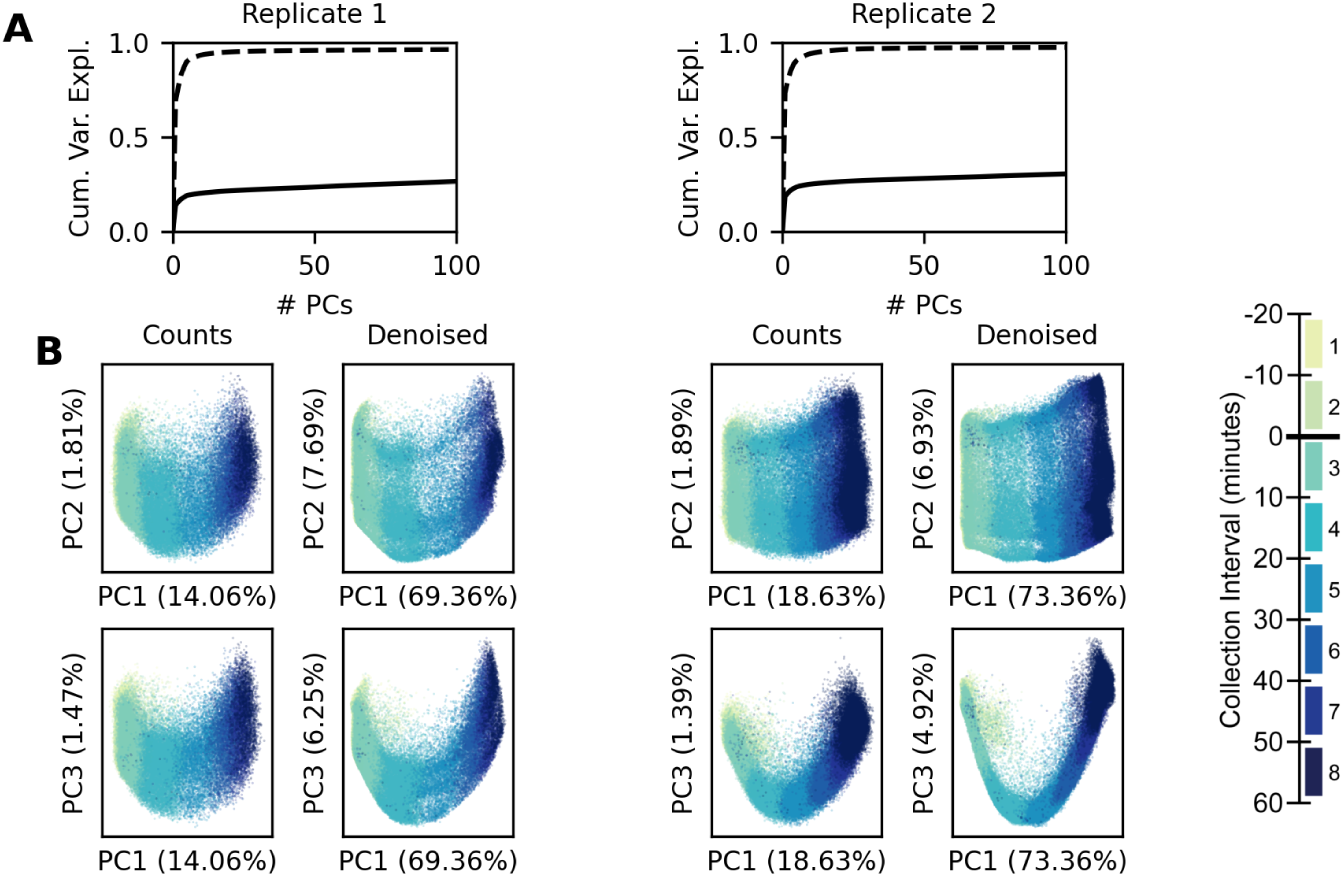
Denoising count data (A) Cumulative variance explained per principal component for raw count data (solid line) and denoised count data (dashed line), plotted separately for each experimental replicate (B) Principal components PC1 & PC2 and PC1 & PC3 plotted against each other for raw count data and denoised count data. Experimental replicates are plotted separately. Individual cells are colored by collection interval.

**Supplemental Figure 18:**
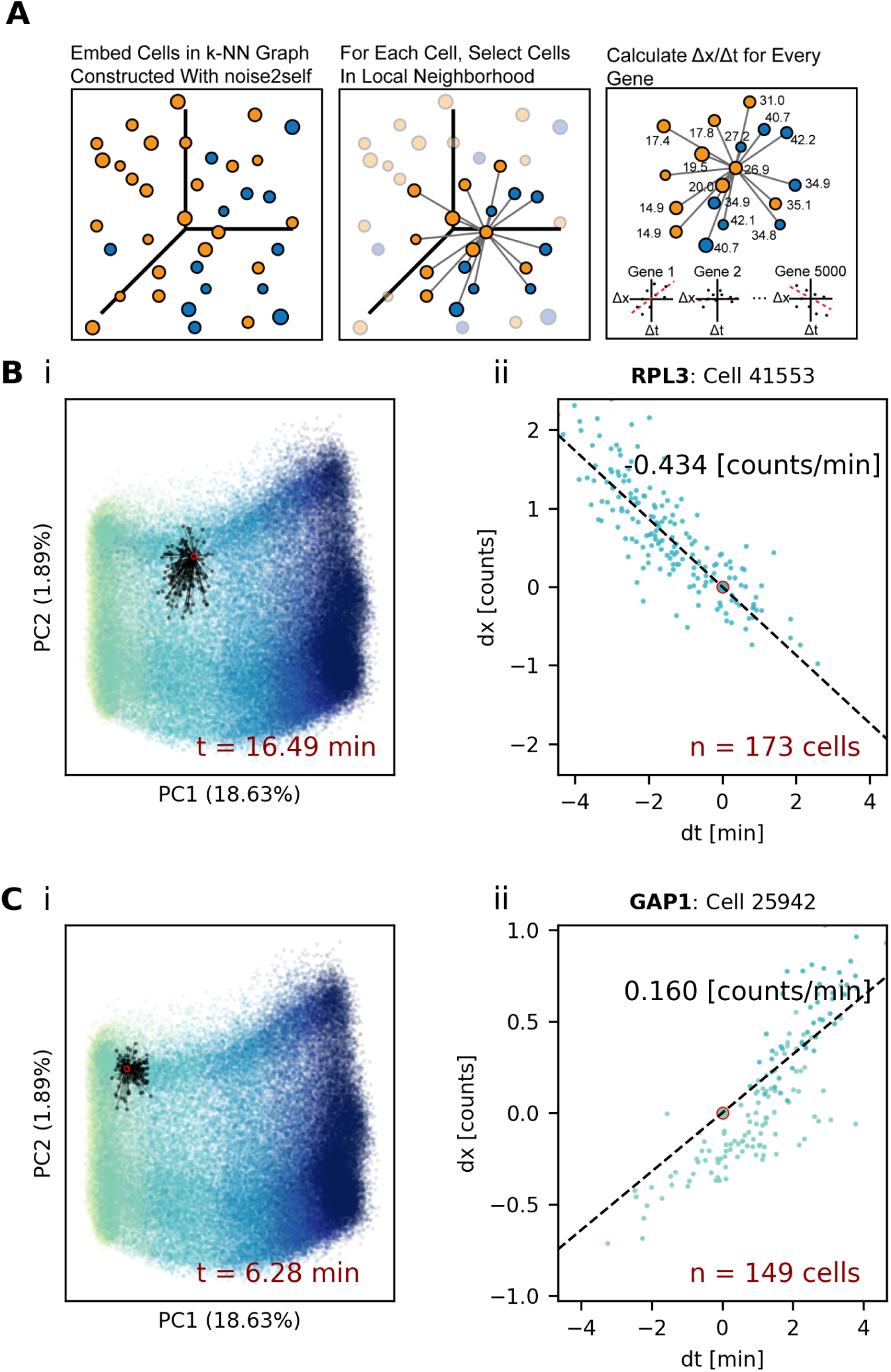
Calculation of RNA velocity (rate of change) (**A**) Schematic showing *k*-NN network construction and RNA velocity regression within local neighborhood for each cell. (**B**) RNA velocity regression for ***RPL3*** at a randomly chosen cell, highlighted on the PC plot (i) and showing only cells connected in the *k*-NN network for velocity regression. (**C**) RNA velocity regression for ***GAP1*** at a random_69_ly chosen cell as in **B**.

**Supplemental Figure 19:**
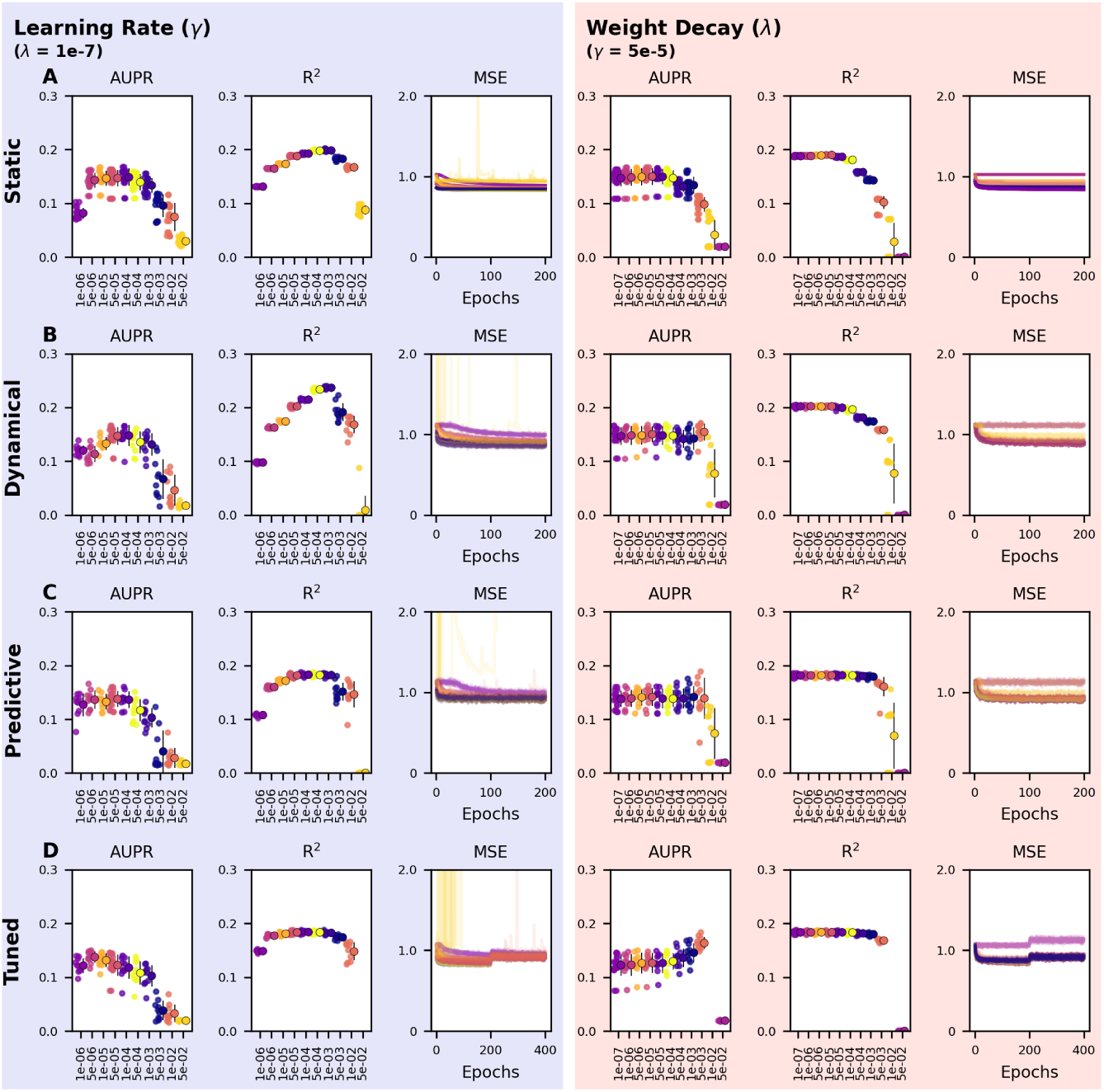
Hyperparameter search for rapamycin velocity inference models, with cells binned as in Supplemental Figure 15 Models are trained with counts as input and RNA velocity as output. (**A**) Static model performance quantified by Area Under the Precision Recall curve (AUPR), coefficient of determination (R^2^), and mean squared error (MSE) per training epoch. Performance is shown for a range of Adam optimizer learning rates (*γ*) with weight decay (*λ*) held constant, and a range of weight decays with learning rate held constant. (B) Dynamical model (*t* + 1 prediction) performance quantified as in A (C) Predictive model (10 sequential predictions) performance quantified as in A. Model input at *t* + 1 is counts *t* plus velocity *t*. (D) Tuned model (training 200 epochs with the dynamical training strategy, followed by 200 epochs with the predictive model training strategy) performance quantified as in A

**Supplemental Figure 20:**
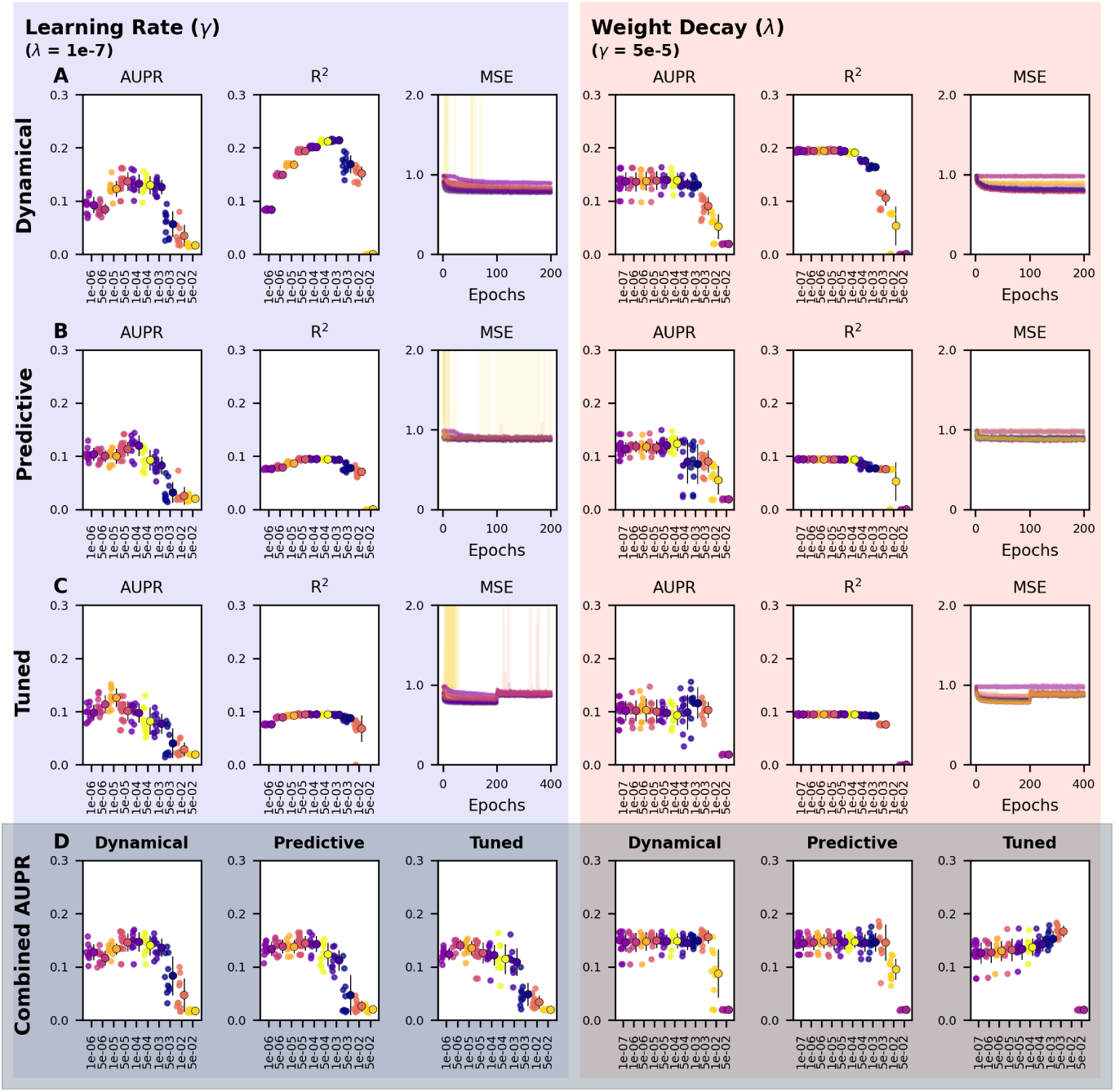
Hyperparameter search for cell cycle velocity models, with cells binned as in Supplemental Figure 16 Models are trained with counts as input and RNA velocity as output. (**A**) Dynamical model (*t* + 1 prediction) performance quantified by Area Under the Precision Recall curve (AUPR), coefficient of determination (R^2^), and mean squared error (MSE) per training epoch. Performance is shown for a range of Adam optimizer learning rates (*γ*) with weight decay (*λ*) held constant, and a range of weight decays with learning rate held constant. (**B**) Predictive model (10 sequential predictions) performance quantified as in **A**. Model input at *t* + 1 is counts *t* plus velocity *t*. (**C**) Tuned model (training 200 epochs with the dynamical training strategy, followed by 200 epochs with the predictive model training strategy) performance quantified as in **A**. (**D**) Regulatory networks combined from rapamycin and cell cycle velocity models by taking the maximum explained relative variance for all regulatory edges from each model. Performance is evaluated by AUPR.

**Supplemental Figure 21:**
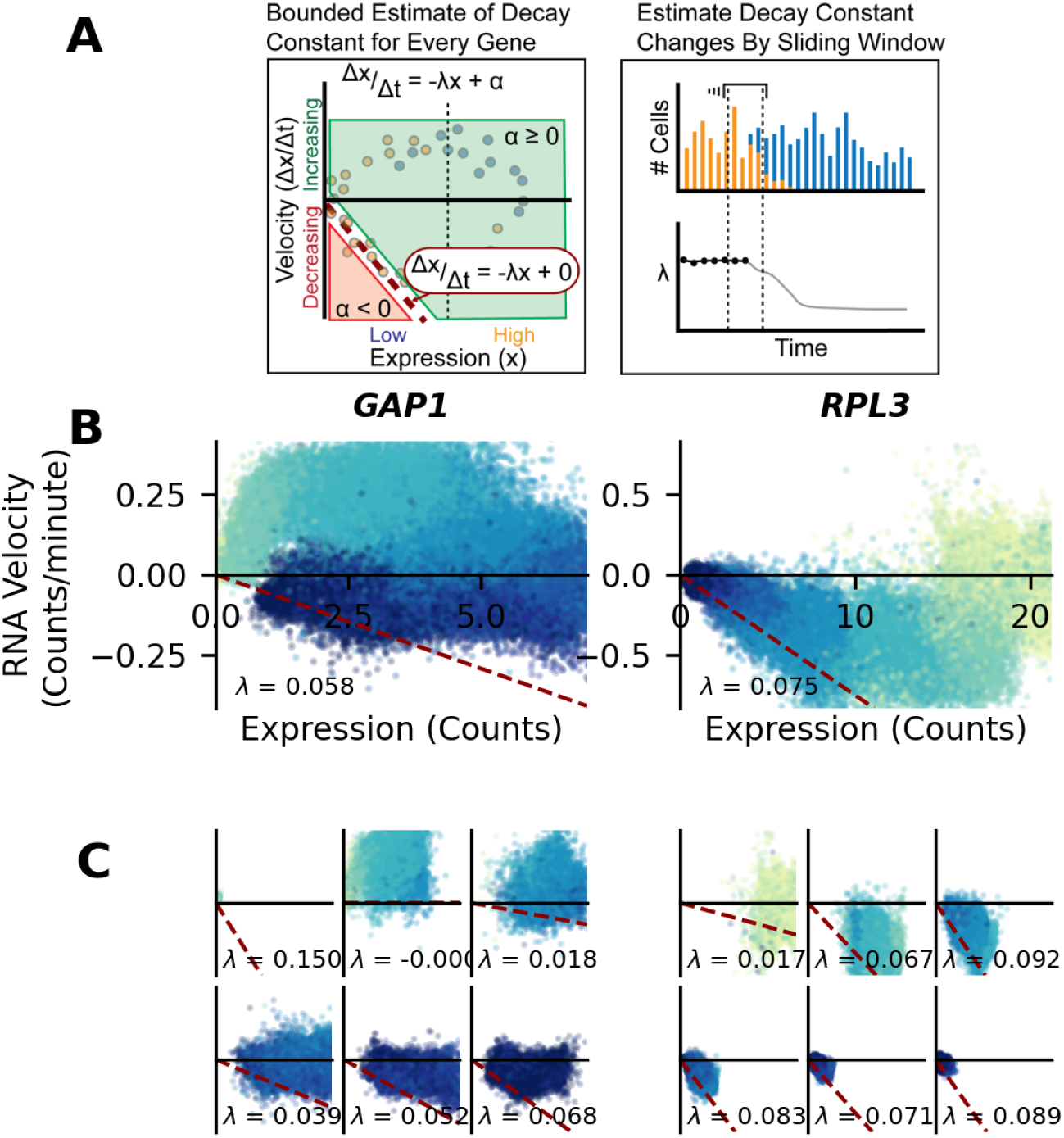
Calculation of RNA decay rates as a dynamic, time-dependent parameter (**A**) Schematic showing bounded estimate of decay rates (**B**) Bounded estimate of *GAP1* and *RPL3* decay constant for all cells (**C**) Bounded estimate of decay rates from 10 minutes of cells (for each gene, -10 to 0, 0 to 10, 10 to 20 left to right in the top row; 20 to 30, 30 to 40, 40 to 50 left to right in the bottom row)

**Supplemental Figure 22:**
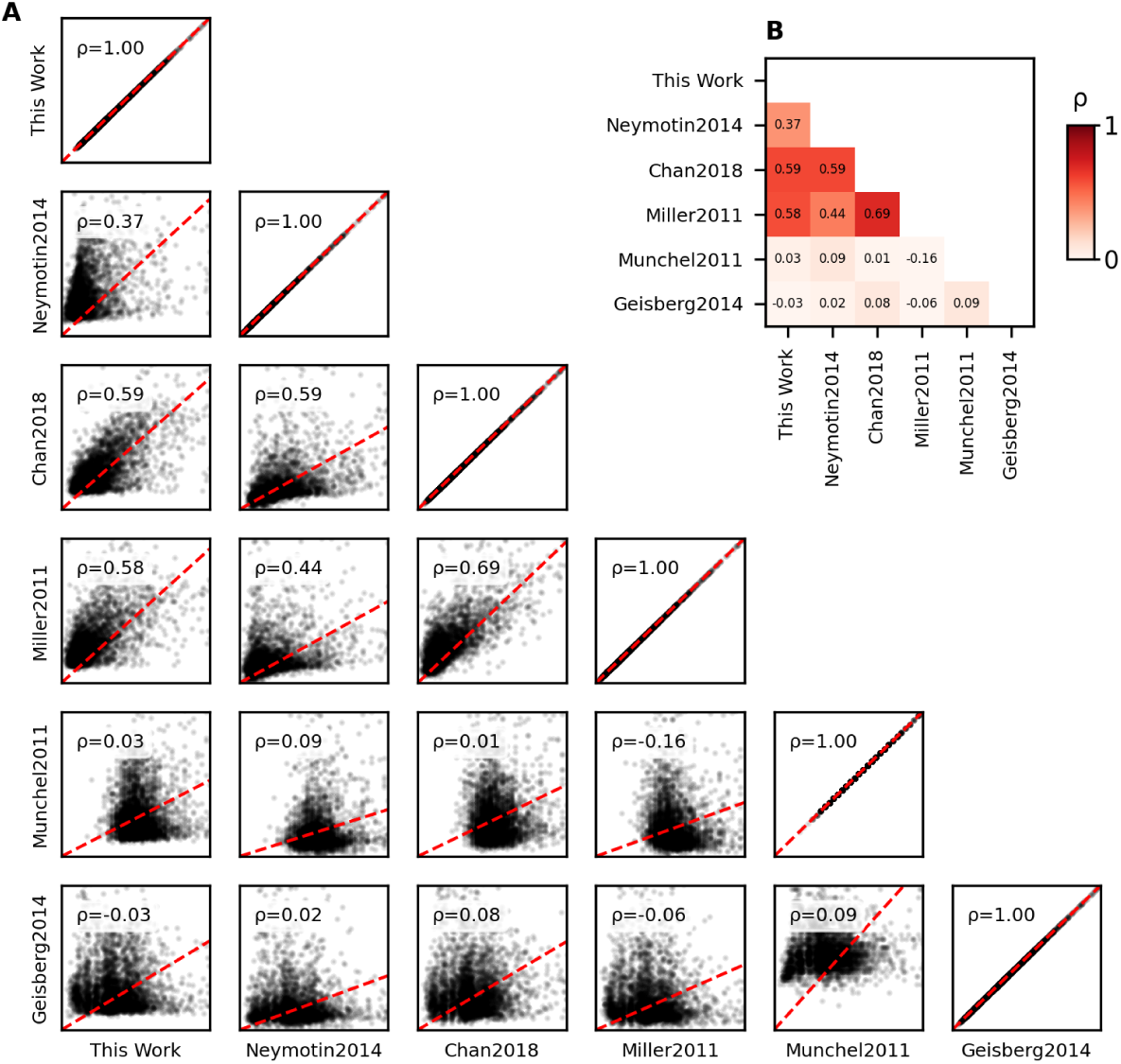
Comparison of estimated mRNA half-lives prior to rapamycin treatment (This Work) with published mRNA half-lives measured by 4-thiouracil labeling in Neymotin2014 (*69*), Chan2018 (*70*), Munchel2011 (*36*), and Miller2011 (*71*). Also compared to decay rates measured by RNA polymerase anchor-away in Geisberg2014 (*72*). (**A**) Half-lives plotted for each data set. Dashed red line is linear regression with no intercept, and fit is reported as spearman correlation coefficient. (**B**) Heatmap of correlation coefficient from **A**.

**Supplemental Figure 23:**
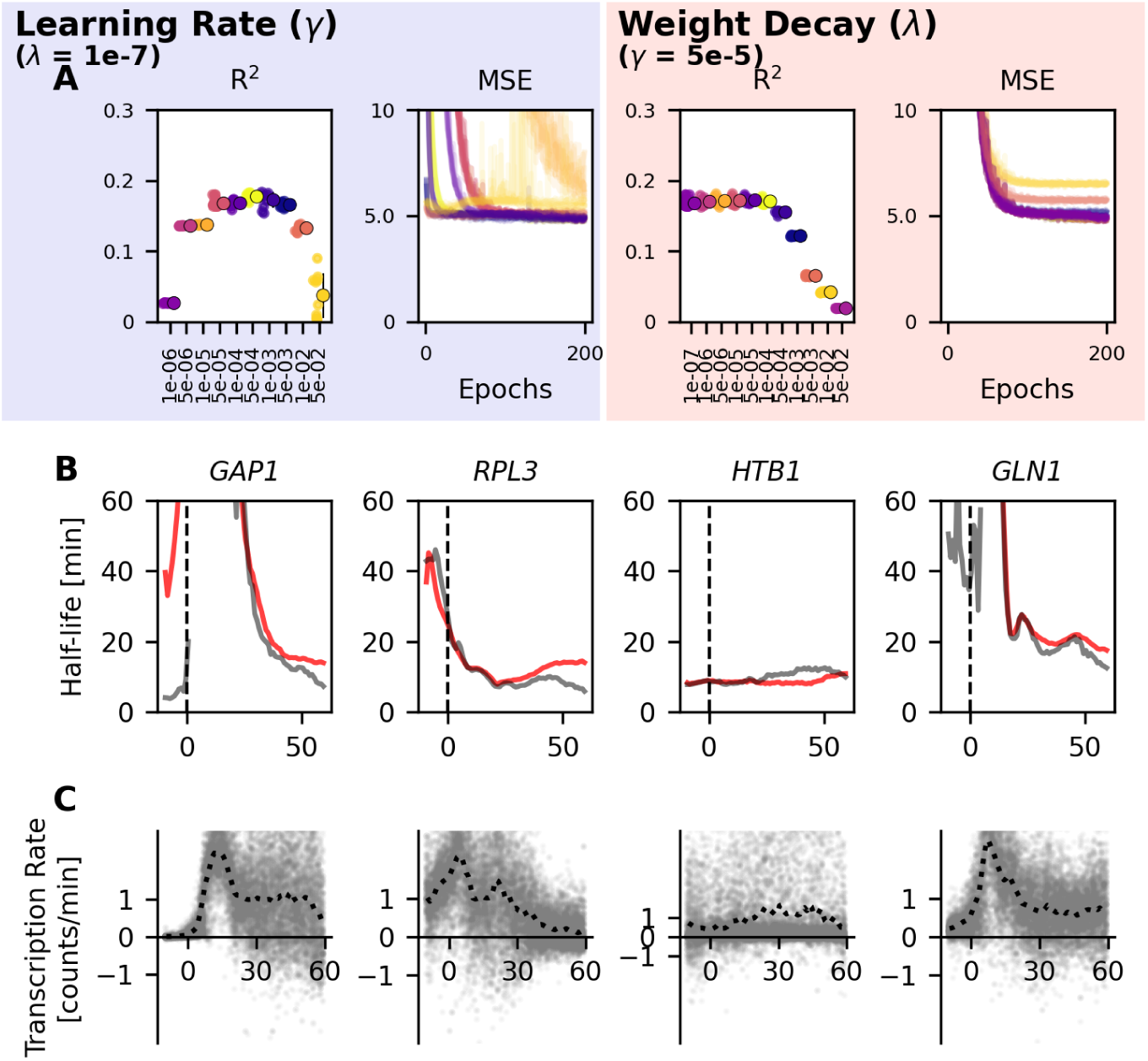
Hyperparameter search for RNA decay model, with cells binned as in Supplemental Figure 15. Models are trained with counts as input and inferred decay rate as output. (**A**) Dynamical model (*t* + 1) performance quantified by coefficient of determination (R^2^), and mean squared error (MSE) per training epoch. Performance is shown for a range of Adam optimizer learning rates (*γ*) with weight decay (*λ*) held constant, and a range of weight decays with learning rate held constant. (**B**) RNA half-life estimates from trained decay model for *GAP1*, *RPL3*, *HTB1*, and *GLN1* (red lines), compared to the RNA half-life estimates used as training data (gray). Training data RNA half-life estimates were calculated as in Supplemental Figure 21, using one-minute sliding windows. (**C**) RNA transcriptional rate estimates determined by subtracting estimated decay velocity (*λX*), with *λ* determined in **B**, from mRNA velocity

**Supplemental Figure 24:**
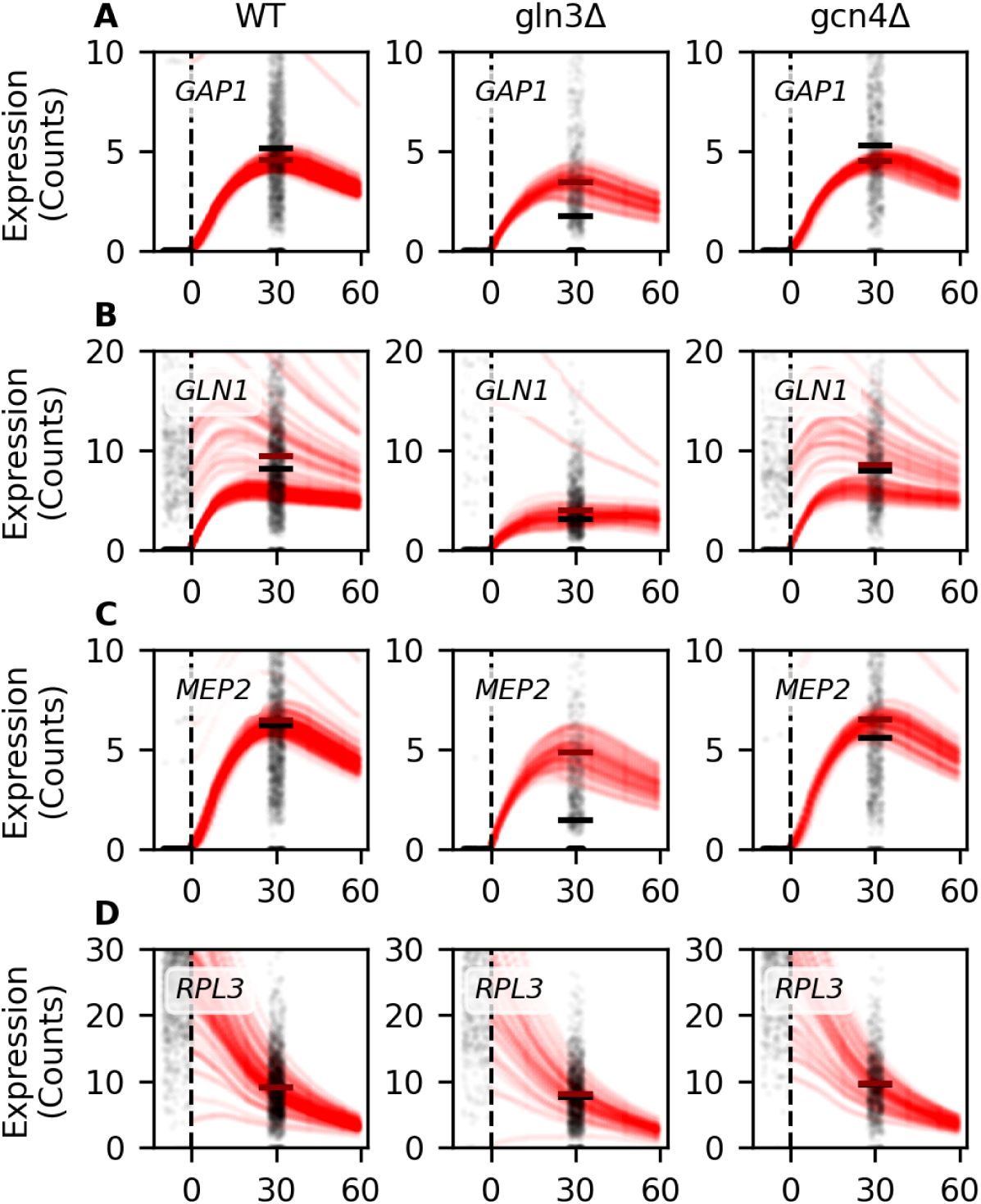
Rapamycin response transcriptional predictions for cells with TFs deleted. *In silico* TF deletion sets TF activity to 0 at all times during prediction. (**A**) Model predictions for *GAP1* in an unperturbed wild-type model, a gln3Δ model, and a gcn4Δ model. Black points are observed counts from untreated (t=0 minutes) and treated (t=30 minutes) cells of the appropriate genotype, jittered for display. Red points are model-predicted counts. Thick black line at t=30 represents the mean of observed data. Thick red line at t=30 represents the mean of all predicted values between 28-32 minutes. (**B**) Model predictions for *GLN1*, plotted as in **A** (**C**) Model predictions for *MEP2*, plotted as in **A** (**D**) Model predictions for *RPL3*, plotted as in **A**

